# Longitudinal examination of cerebellar morphometry across typical aging adults reveals nonlinear heterogenous trends

**DOI:** 10.64898/2026.11.13.766545

**Authors:** Zachary M. Brodnick, Naomi Doron, Jessica A. Turner

## Abstract

Cerebellar structural integrity has been associated with cognitive capability in both neurodegenerative and healthy aging populations. The understanding of regional, longitudinal cerebellar structural changes is not well studied, and with an increasing aging population, there is an increasing need to fill this gap. Moreover, traditional modeling approaches apply linear techniques, which may fail to detect possible nuanced aging patterns. The present study investigates the nonlinear effect of age on subregional cerebellar structures in healthy adults (n = 1,371), aged 36 to 90+, from the Human Connectome Project in Aging and the follow-up continuation, Aging Adult Brain Connectome studies. Subject T1w images underwent cerebellar segmentation and volumetric estimation via the Automatic Cerebellum Parcellation using U-NET with Locally Constrained Optimization (ACAPULCO) algorithm. Cross-sectional, longitudinal, and sex specific volumetric differences were examined using generalized additive mixed models. Heterogenous atrophy patterns were observed across the cerebellum; however, the majority of effects were evident in the posterior cerebellum. Longitudinal models showed nonlinear negative trends for all regions surviving correction, while baseline age effects had a combination of linear and nonlinear negative trajectories after correction. Specific sex differences emerged, where men predominantly had higher regional volumes, as well as there being sex-related influences of longitudinal morphologic patterns. Furthermore, midlife appeared as a period of structural vulnerability, demonstrating escalated volume loss. Consistent with the literature, the identified volumetric reduction patterns suggest that age effects on the cerebellum are multifaceted, with region specific alterations, highlighting the complexity of this structure.

## Introduction

Aging is a natural biological transition period for humans, with projected rates of older populations growing coinciding with increased life expectancy (Caplan, 2023; World Health Organization (WHO), 2025). This period of life may be marred by motor and subtle cognitive decline (Harada et al., 2013), coupled with neurobiological alterations (Cabeza et al., 2002; Luo et al., 2019; Setton et al., 2022). A brain region gaining increased recognition as a key area involved in age-related changes is the cerebellum. Emerging evidence highlights the importance of cerebellar structural integrity in healthy aging; however, regional and longitudinal trends remain poorly understood.

The study of the cerebellum, historically defined as a motor region, has undergone a renaissance, as the structure has been increasingly linked to cognitive and affective processing (Schmahmann, 1996; Stoodley & Schmahmann, 2009; Stoodley, 2012; Guell et al., 2018). A recent rhesus monkey lesion study has shown a causal link between deep cerebellar nuclei insults and working memory and executive functioning impairment (Schmahmann et al., 2025), suggesting the findings from a meta-analysis of human cerebellar neuromodulation by Gatti et al. (2021) extend across evolutionarily related species. In the context of human aging, the cerebellum may be a modulatory reserve region involved in cerebral compensation and refinement of cognitive processes (Devita et al., 2024; Mitoma et al., 2020). Empirical evidence focusing on structural MRI in aging populations highlights this (Bernard & Seidler 2013; Cooper et al., 2023; Hicks et al., 2024; d’Oleire Uquillas et al., 2026), suggesting that cerebellar structural integrity may be necessary for neurocognitive performance. Therefore, studying patterns of structural decline in the cerebellum may provide further insight into neurodegeneration and cognitive decline in the aging brain.

Across development, the cerebellum has been found to exhibit non-linear patterns. Several studies have consistently shown that posterior cerebellar regions follow an inverted-U maturation pattern, with areas peaking in adolescence or early adulthood (Bernard et al., 2014, Gil-Paterna et al., 2026, Romero et al., 2021; Tiemeier et al., 2010). Likewise, nonlinear techniques like quadratic, logarithmic, and polynomial modeling have successfully been applied in older adult samples, suggesting negative age effects (Bernard et al., 2014; Hicks et al., 2022; Romero et al., 2021; Wang et al., 2024; Ziegler et al., 2011). Moreover, it has been observed that regional volumetric patterns may exist. One study found that posterior regions exhibit inverted quadratic trends, Crus I followed a negative linear trend, while vermal and anterior regions followed logarithmic trends across age (Bernard et al., 2014). Further support for this divergent declining pattern was observed in older adults. In particular, Han et al. (2020a) and Romero et al. (2021) both found that anterior lobe regions exhibited descending curved trends. However, Han et al. (2020a) showed posterior lobe regions declined linearly, and Romero et al. (2021) reported additional nonlinear negative trends.

Generalized additive mixed models (GAMMs) are well positioned to explore these discrepant findings. GAMMs allow for flexible modeling of data that otherwise may not be linear as well as data that adhere to heterogeneous trajectories across time. In doing so, they enable the exploration of complex trends while avoiding restrictive prior assumptions that could potentially mask meaningful results. Thus, they are a recommended technique to accurately examine lifespan variations (Hastie & Tibshirani, 1986; Sørensen et al., 2021, Wood, 2017) and have been increasingly used in both neurocognitive and structural lifespan studies, in healthy and clinical populations (Bethlehem et al., 2022; Lammer et al., 2025; Herting et al., 2018; Jones et al., 2023; Poortman et al., 2024; Schuff et al., 2012).

While several studies have investigated longitudinal trends (Tiemierer 2010, Han et al 2020a, Cooper et al. 2023; Raz et al., 2003; 2005; 2010; 2013), only Han et al. (2020a) investigated the cerebellum in an in-depth manner across regions. There has been a rise of detailed parcellation studies modeling age differences cross-sectionally (Bernard et al., 2014, Cui et al., 2020; d’Oleire Uquillas et al., 2026; Gil-Paterna et al., 2026, Hicks et al., 2022, Koppelmans et al., 2017; Steele & Chakravarty, 2018; Wang et al., 2024). To accurately assess within-subject changes that are undetected in cross-sectional studies, longitudinal approaches are necessary. This is further complemented by the notion that there are exceptions to cerebellar organization uniformity; studies have shown subregional anatomical, molecular, physiological, and functional specificity, underscoring the need to conduct more localized analyses (Cerminara et al., 2015; Nettekoven et al., 2024; Zhou et al., 2014). For an area that houses an estimated 80% of neurons (Sereno et al., 2020), cross-sectional studies may not be poised to adequately assess cerebellar complexity, so that masking of subtle findings and overgeneralizations could ensue.

This study aims to address this gap by exploring cerebellar lobular structural volumes longitudinally through non-linear modeling. Within a healthy aging sample, we investigate both between and within subject volumetric trends with age. We hypothesized that healthy older adults would exhibit lower cerebellar lobular and subregion volumes compared to younger healthy adults. Additionally, we predict that there will be heterogenous negative age effects across cerebellar regions, emphasized in posterior regions. Furthermore, we expect sex differences in volume across age as well as sex modulating longitudinal volumetric trends, as seen in previous work (Han et al., 2020a). Through this work, advancements in what constitutes normative cerebellar age trends could be gained, laying the foundation for future research on cerebellar involvement in aberrant age pathology.

## Methods and materials

### Participants

Participants were recruited from the Human Connectome Project in Aging (HCP-A) and a longitudinal extension study, Aging Adult Brain Connectome (AABC) Study Release 2.0 (Bookheimer et al., 2019; Harms et al., 2018). Subjects (n = 1,396) were cognitively healthy aging adults aged from 36 years to 90+ and underwent up to four timepoints of MRI visits. Filtering out subjects who had at least one structural scan and passed quality assurance (n = 1,371), follow-up intervals were as follows: Visit 2 (*M* = 3.0 years from baseline, SD =1.7); Visit 3 (*M* = 5.6 years from baseline, *SD* = 1.0); Visit 4 (*M* = 6.9 years from baseline, *SD* = 0.7). Of note, disruptions from the COVID-19 pandemic contributed to some of the variability in follow-up intervals, particularly at Visit 2. Additional information on recruitment strategies, cognitive capacity assessment, and inclusion and exclusion criteria can be found elsewhere (Bookheimer et al., 2019). Sample demographics for those who passed quality assurance, details later discussed in the methods, are summarized in **Table 1**. Of note, subjects who were 90 years or older were coded as “90 or older” via the AABC protocol, therefore in the present study they were assigned a binned age of 90+. Follow-up intervals and study site breakdown are summarized in **Table 2**. Blood pressure classification was based on recent American Heart Association recommendations (American Heart Association, 2025). Further details about the temporal stability of subject blood pressure classification, see **Supplemental Fig. 1.**

**Table 1.** Sample Baseline Demographics.

|  | **36–55<br>(N=564)** | **56-69<br>(N=335)** | **70–90+<br>(N=472)** | **Overall<br>(N=1371)** |
|---|---|---|---|---|
| **Age (years)** |  |  |  |  |
| Mean (SD) | 45.6 (5.98) | 62.4 (3.93) | 80.0 (6.40) | 61.5 (16.0) |
| Median [Min, Max] | 46.0 [36.0, 55.0] | 62.0 [56.0, 69.0] | 80.0 [70.0, 90.0] | 61.0 [36.0, 90.0] |
| **Sex** |  |  |  |  |
| F | 334 (59.2%) | 197 (58.8%) | 251 (53.2%) | 782 (57.0%) |
| M | 230 (40.8%) | 138 (41.2%) | 221 (46.8%) | 589 (43.0%) |
| **Race** |  |  |  |  |
| American Indian/Alaska Native | 1 (0.2%) | 1 (0.3%) | 2 (0.4%) | 4 (0.3%) |
| Asian | 54 (9.6%) | 24 (7.2%) | 9 (1.9%) | 87 (6.3%) |
| Black or African American | 139 (24.6%) | 65 (19.4%) | 59 (12.5%) | 263 (19.2%) |
| Hawaiian or Pacific Islander | 3 (0.5%) | 1 (0.3%) | 0 (0%) | 4 (0.3%) |
| More than one race | 41 (7.3%) | 9 (2.7%) | 5 (1.1%) | 55 (4.0%) |
| White | 299 (53.0%) | 233 (69.6%) | 396 (83.9%) | 928 (67.7%) |
| Missing | 27 (4.8%) | 2 (0.6%) | 1 (0.2%) | 30 (2.2%) |
| **Ethnicity** |  |  |  |  |
| Hispanic or Latino | 118 (20.9%) | 29 (8.7%) | 23 (4.9%) | 170 (12.4%) |
| Not Hispanic or Latino | 444 (78.7%) | 305 (91.0%) | 447 (94.7%) | 1196 (87.2%) |
| Missing | 2 (0.4%) | 1 (0.3%) | 2 (0.4%) | 5 (0.4%) |
| **Education** |  |  |  |  |
| Less than high school | 107 (19.0%) | 62 (18.5%) | 59 (12.5%) | 228 (16.6%) |
| High school graduate | 41 (7.3%) | 18 (5.4%) | 45 (9.5%) | 104 (7.6%) |
| Some college | 114 (20.2%) | 60 (17.9%) | 89 (18.9%) | 263 (19.2%) |
| College graduate | 155 (27.5%) | 103 (30.7%) | 113 (23.9%) | 371 (27.1%) |
| Graduate degree | 145 (25.7%) | 90 (26.9%) | 166 (35.2%) | 401 (29.2%) |
| Missing | 2 (0.4%) | 2 (0.6%) | 0 (0%) | 4 (0.3%) |
| **MoCA Total** |  |  |  |  |
| Mean (SD) | 26.5 (2.46) | 26.4 (2.37) | 24.9 (2.80) | 26.0 (2.66) |
| Median [Min, Max] | 27.0 [20.0, 30.0] | 27.0 [19.0, 30.0] | 25.0 [17.0, 30.0] | 26.0 [17.0, 30.0] |
| Missing | 1 (0.2%) | 0 (0%) | 1 (0.2%) | 2 (0.1%) |
| **Blood Pressure Classification** |  |  |  |  |
| Normal | 182 (32.3%) | 67 (20.0%) | 61 (12.9%) | 310 (22.6%) |
| Elevated | 63 (11.2%) | 37 (11.0%) | 60 (12.7%) | 160 (11.7%) |
| Stage 1 | 193 (34.2%) | 100 (29.9%) | 113 (23.9%) | 406 (29.6%) |
| Stage 2 | 120 (21.3%) | 121 (36.1%) | 227 (48.1%) | 468 (34.1%) |
| Severe | 1 (0.2%) | 4 (1.2%) | 4 (0.8%) | 9 (0.7%) |
| Missing | 5 (0.9%) | 6 (1.8%) | 7 (1.5%) | 18 (1.3%) |

*Note.* Demographics reflect participants who passed QC and were used in analyses. Information was grouped via pragmatically defined age groups for ease of visualization. For Blood Pressure Classification, Stage 1, Stage 2, and Severe denote hypertension stages. SD = standard deviation.

**Table 2.** Number of Participants at Each Study Visit and Site.

| **Visit** | **Total** | **MGH** | **UCLA** | **UMinn** | **WashU** |
|---|---|---|---|---|---|
| V1 | 1,367 | 302 | 322 | 326 | 417 |
| V2 | 857 | 218 | 193 | 223 | 223 |
| V3 | 431 | 109 | 69 | 134 | 119 |
| V4 | 86 | 33 | 8 | 18 | 27 |

*Note.* Counts reflect unique participants with MRI data at each visit who passed QC.

### MRI Acquisition and Processing

Scanning occurred across four sites, where all adhered to the Human Connectome Project (HCP) protocol (Harms et al., 2018). Structural MPRAGE T1-weighted images acquired at each visit used a 3-Tesla Siemens Prisma, with a 32-channel head coil. Specific scanner parameters include: voxel size = 0.8 mm3, FOV = 256 × 240 × 166, flip angle = 8 degrees, TR/TI = 2500/1000 ms, Multiecho TE = 1.8/3.6/5.4/7.2 ms. Furthermore, T1-weighted images used for analyses underwent the HCP minimal preprocessing pipeline, PreFreeSurfer step (Glasser et al., 2013), where bias field and distortion correction as well as ACPC alignment were applied to selected images.

For cerebellar parcellation, the Automatic Cerebellum Parcellation using U-NET with Locally Constrained Optimization (ACAPULCO) algorithm was employed (Han et al., 2020b; Kerestes et al. 2022). In brief, this pipeline isolates the cerebellum into 28 subregions, calculates volume as well as using FreeSurfer Version v8.1.0 (Fischl, 2012) to calculate total intracranial volume (TIV). To use as a later control, TIV was grand mean centered for each subject based on the entire sample TIV average. Similar to Han et al. (2020a), select cerebellar subregions were combined for post hoc analyses following anatomical classification by Schmahmann et al. (2000) to allow for comparisons to previous literature findings. A detailed list of regions of interest (ROIs) can be found in **Table 3**.

**Table 3.**
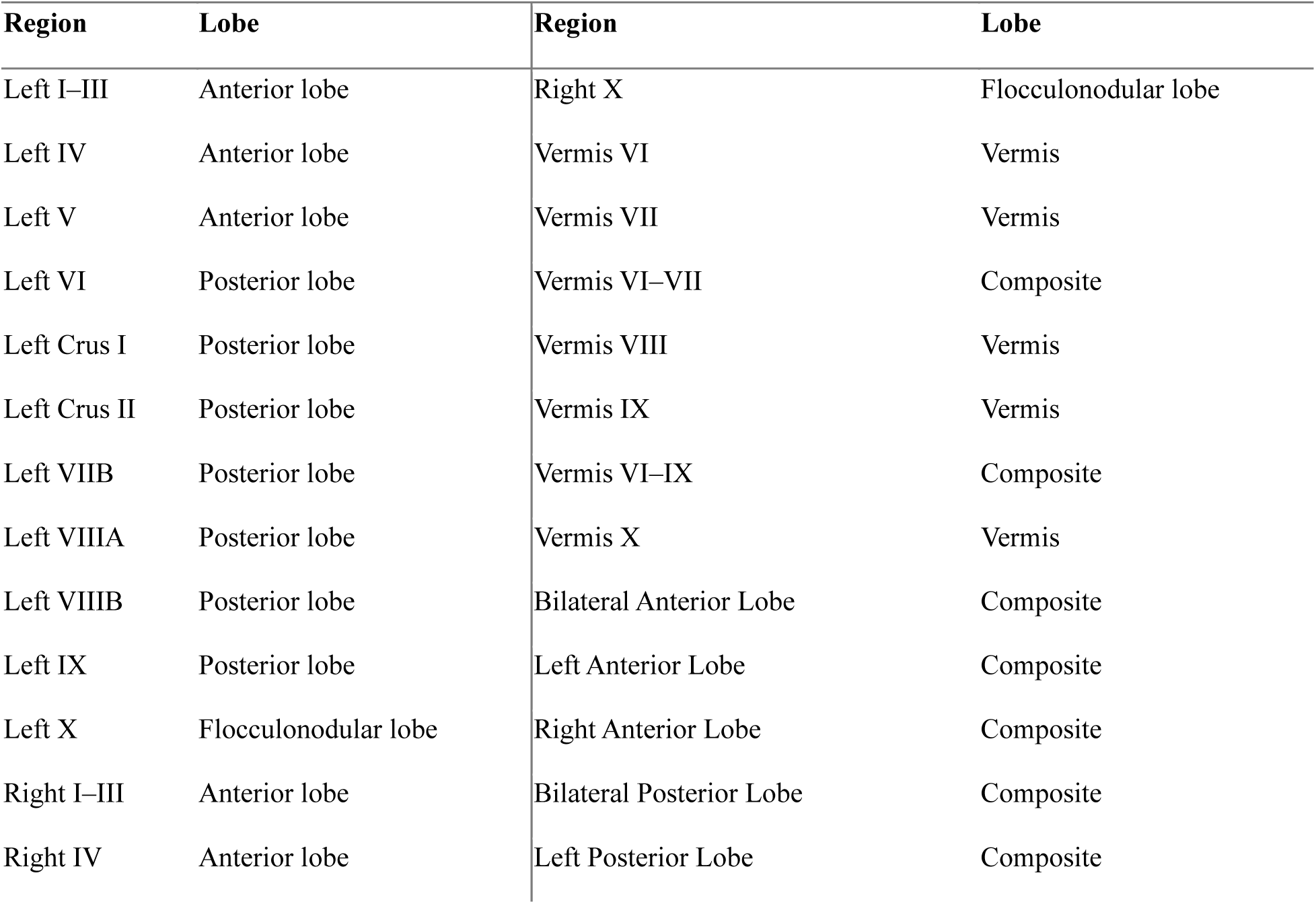
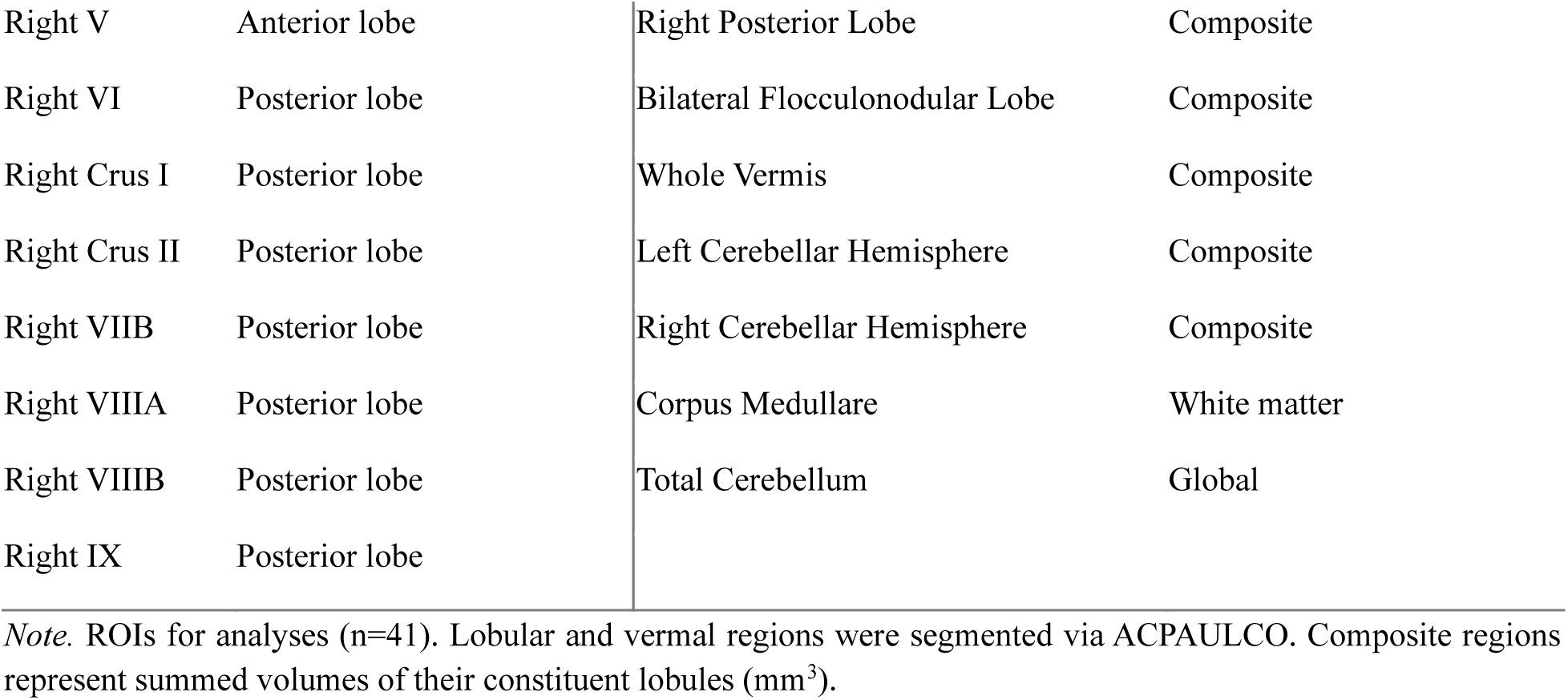
Cerebellar ROIs.

| Region | Lobe | Region | Lobe |
|---|---|---|---|
| Left I–III | Anterior lobe | Right X | Flocculonodular lobe |
| Left IV | Anterior lobe | Vermis VI | Vermis |
| Left V | Anterior lobe | Vermis VII | Vermis |
| Left VI | Posterior lobe | Vermis VI–VII | Composite |
| Left Crus I | Posterior lobe | Vermis VIII | Vermis |
| Left Crus II | Posterior lobe | Vermis IX | Vermis |
| Left VIIB | Posterior lobe | Vermis VI–IX | Composite |
| Left VIIIA | Posterior lobe | Vermis X | Vermis |
| Left VIIIB | Posterior lobe | Bilateral Anterior Lobe | Composite |
| Left IX | Posterior lobe | Left Anterior Lobe | Composite |
| Left X | Flocculonodular lobe | Right Anterior Lobe | Composite |
| Right I–III | Anterior lobe | Bilateral Posterior Lobe | Composite |
| Right IV | Anterior lobe | Left Posterior Lobe | Composite |
| Right V | Anterior lobe | Right Posterior Lobe | Composite |
| Right VI | Posterior lobe | Bilateral Flocculonodular Lobe | Composite |
| Right Crus I | Posterior lobe | Whole Vermis | Composite |
| Right Crus II | Posterior lobe | Left Cerebellar Hemisphere | Composite |
| Right VIIB | Posterior lobe | Right Cerebellar Hemisphere | Composite |
| Right VIIIA | Posterior lobe | Corpus Medullare | White matter |
| Right VIIIB | Posterior lobe | Total Cerebellum | Global |
| Right IX | Posterior lobe |  |  |

*Note.* ROIs for analyses (n=41). Lobular and vermal regions were segmented via ACPAULCO. Composite regions represent summed volumes of their constituent lobules (mm<sup>3</sup>).

ACAPULCO output was assessed through two quality assurance streams. Automated outlier detection via the ENIGMA Cerebellum Volumetrics Pipeline, as well as manual visual inspection of cerebellar segmentation to evaluate accuracy of cerebellar coverage and quality were utilized. In total, 47 whole scans were excluded (27 failed segmentation and 20 image quality). Additionally, 713 subjects had at least one lobule excluded with a mean of 1.96 lobules being excluded out of that subgroup. An important caveat is that no subjects were identified to have ever received a Mild Cognitive Impairment (MCI) or Alzheimer’s (AD) diagnosis at baseline; however, 8 subjects later reported receiving a diagnosis (n = 2 for AD, n = 6 for MCI). Regardless, these subjects passed quality assurance and hence were included in subsequent analyses.

### Statistical Analysis

To model the effect of age on cerebellar subregional volumes, generalized additive models (GAMs) and GAMMs were utilized in R (v 4.5.2) (R Core Team, 2025). Briefly, GAMs are data driven extensions of regressions that employ smooth spline functions on predictors to model nonlinear relationships, without priori assumptions of linearity (Hastie & Tibshirani, 1986; Wood, 2017). Likewise, GAMMs are hierarchical iterations of GAMs, whereby they are equipped to model random effects as well as cross-sectional and longitudinal effects using the same framework as linear mixed-effects models (LMMs) (Wood, 2017). Notably, because both modeling techniques introduce smooth splines, no single slope for age can be derived. First derivative plots address this issue by calculating the instantaneous rate of change per age, allowing for the evaluation of both magnitude and direction of smoothed age effects. For more information, see **Supplemental Methods** for details on how first derivatives were computed.

Two models were constructed for comparison, both of which applied penalized smooth terms on age specified by restricted maximum likelihood (REML), as well as using cubic regression splines as the basis function. The first was a cross-sectional model using the GAM framework from the *mgcv* package (v1.9.4), to examine the cross-sectional effects of age across all 41 subregions. Here, only baseline data were selected, and we introduced a smooth term on age, while controlling for mean centered TIV, sex, and random scanner effects. The motivation for the cross-sectional analysis was to provide a basis for comparison with recent cross-sectional work (d’Oleire Uquillas et al., 2026; Hicks et al., 2022; Wang et al., 2024). Subsequent follow-up sensitivity analyses were run, introducing control variables such as blood pressure classification, race, ethnicity, and education history.

Extending this approach, due an absence of cohort effects in the dataset, from the recommendation of Sørensen et al. (2021), the second model examined the nonlinear effect of age at each timepoint using GAMMs formulation via the *gamm4* package (v0.3.0) on all 41 subregions. An identical procedure was adopted for this longitudinal model from the cross-sectional model, however with visit specific age and TIV predictors, along with the addition of random subject and scanner intercepts. Additionally, a third separate model was constructed to examine the interaction of age and sex across time. Following the same format as the longitudinal model, a smoothed factor interaction between sex and age using up to five basis functions was added.

The results of all models underwent correction for multiple comparisons across the cerebellar regions using Benjamini-Hochberg false discovery rate (FDR) (Benjamini & Hochberg, 1995) (p < 0.05). Additionally, separate sensitivity analyses were run, controlling for the fixed effects of blood pressure classification, education, race, ethnicity, and cognitive function via the Montreal Cognitive Assessment (MoCA) (Nasreddine et al., 2005) to assess the survival of age effects.

## Results

### Cross-Sectional Model

Significant age effects surviving FDR correction were revealed for 34 of the 41 ROIs for the cross-sectional analyses. Of those significant regions, 25 showed nonlinear trends. Additionally, all significant ROIs showed negative age effects, indicating that older adults have smaller volumes than younger adults. Furthermore, baseline sex differences were detected, with males predominantly exhibiting larger volumes in select regions. See **Table 4** below for a breakdown of the fixed and smooth effects per region and **Fig. 1** for example first derivative plots (remaining ROI first derivative plots can be found in **Supplemental Fig. 2**).

**Fig 1.**
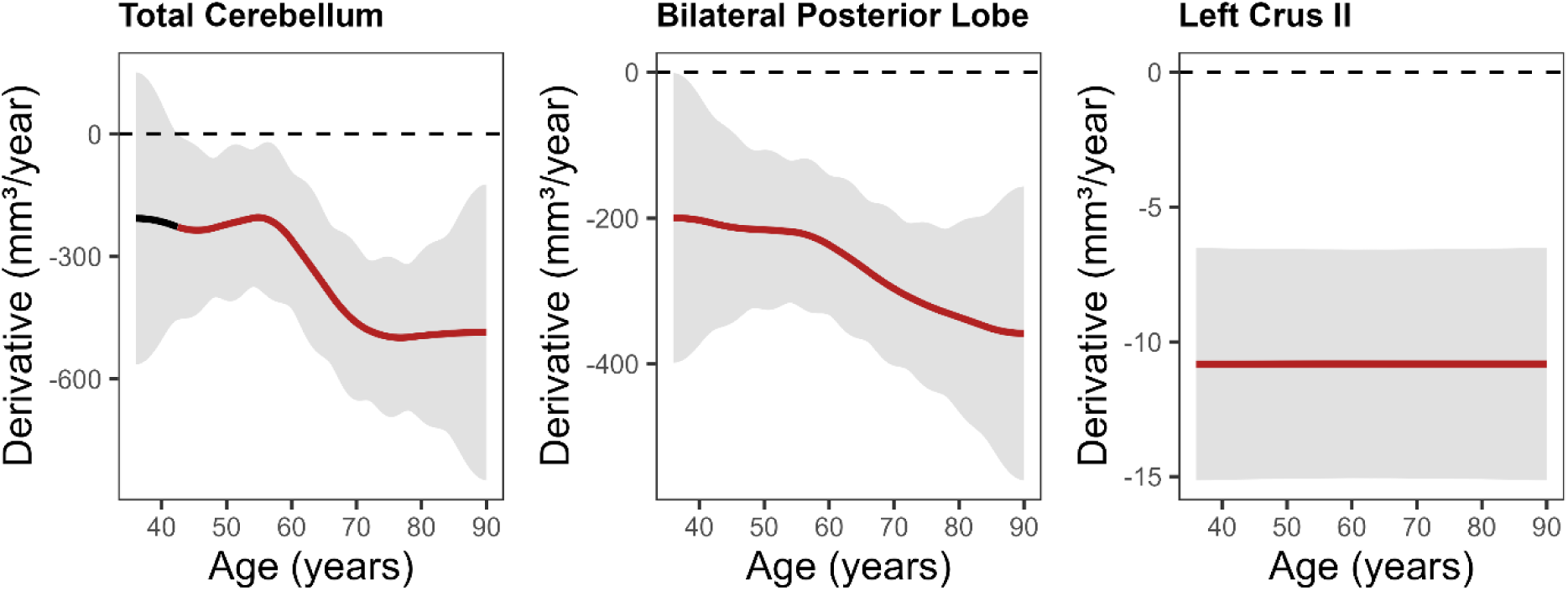
Panel of first derivative plots of smoothed age effects from the baseline models for Total Cerebellum, Bilateral Posterior Cerebellum, and Left Crus II volume, evaluating ages where the instantaneous rate of volume change (mm³/year) was significantly different from zero. Red segments indicate significance, where the 95% confidence interval grey band does not overlap with zero. The dashed horizontal line marks zero (no change in volume). These results show examples of the acceleration of volume loss between 50-70 years, in total cerebellar volume and the Bilateral Posterior Lobe, compared with the constant volume loss estimated in Left Crus II. The plateauing effect around age 90 may be attributed to the binning of participants who were reported to be 90 years of age or older.

**Table 4.** Sex and Age effects on cerebellar subregional volumes from the cross-sectional model.

|  | ***Sex (Male)*** |  |  |  | ***s(age)*** |  |  |
|---|---|---|---|---|---|---|---|
| **Region** | **Estimate** | ***SE*** | ***t*-value** | ***p*** | ***edf*** | ***F*** | ***p*** |
| Left I-III | 18.67 | 10.57 | 1.77 | 0.078 | 2.28 | 2.56 | 0.045 |
| Left IV | 300.12 | 32.53 | 9.23 | **< .001** | 1.01 | 0.10 | 0.766 |
| Left V | 173.35 | 27.38 | 6.33 | **< .001** | 2.43 | 34.79 | **< .001** |
| Left VI | -339.41 | 84.12 | -4.03 | **< .001** | 3.08 | 41.08 | **< .001** |
| Left Crus I | 190.45 | 114.17 | 1.67 | 0.096 | 2.43 | 89.87 | **< .001** |
| Left Crus II | 287.94 | 81.19 | 3.55 | **< .001** | 1.00 | 29.46 | **< .001** |
| Left VIIB | -225.93 | 67.23 | -3.36 | **< .001** | 1.01 | 263.68 | **< .001** |
| Left VIIIA | 573.99 | 68.42 | 8.39 | **< .001** | 2.43 | 13.42 | **< .001** |
| Left VIIIB | 2.78 | 39.46 | 0.07 | 0.944 | 1.01 | 77.27 | **< .001** |
| Left IX | -59.00 | 42.08 | -1.40 | 0.161 | 1.28 | 41.11 | **< .001** |
| Left X | 21.58 | 4.81 | 4.49 | **< .001** | 2.37 | 2.44 | 0.070 |
| Right I-III | 5.79 | 10.73 | 0.54 | 0.590 | 1.92 | 18.23 | **< .001** |
| Right IV | 291.14 | 33.48 | 8.70 | **< .001** | 2.13 | 1.96 | 0.097 |
| Right V | 119.66 | 29.01 | 4.12 | **< .001** | 4.51 | 38.20 | **< .001** |
| Right VI | -357.21 | 78.29 | -4.56 | **< .001** | 2.58 | 43.85 | **< .001** |
| Right Crus I | 301.33 | 120.13 | 2.51 | **0.012** | 2.47 | 91.18 | **< .001** |
| Right Crus II | 319.09 | 82.66 | 3.86 | **< .001** | 1.74 | 29.70 | **< .001** |
| Right VIIB | -86.33 | 67.61 | -1.28 | 0.202 | 1.01 | 290.27 | **< .001** |
| Right VIIIA | 170.62 | 63.53 | 2.69 | **0.007** | 1.01 | 0.16 | 0.693 |
| Right VIIIB | -1.62 | 37.43 | -0.04 | 0.965 | 1.87 | 67.55 | **< .001** |
| Right IX | -13.85 | 41.42 | -0.33 | 0.738 | 1.01 | 73.18 | **< .001** |
| Right X | 9.53 | 4.96 | 1.92 | 0.055 | 2.48 | 7.46 | **< .001** |
| Vermis VI | 11.39 | 13.65 | 0.83 | 0.404 | 2.68 | 14.33 | **< .001** |
| Vermis VII | 5.34 | 10.37 | 0.51 | 0.607 | 1.64 | 3.05 | 0.045 |
| Vermis VI-VII | 16.04 | 19.58 | 0.82 | 0.413 | 2.85 | 11.09 | **< .001** |
| Vermis VIII | 111.24 | 19.77 | 5.63 | **< .001** | 1.01 | 1.92 | 0.165 |
| Vermis IX | -36.14 | 9.34 | -3.87 | **< .001** | 1.68 | 9.98 | **< .001** |
| Vermis VI-IX | 86.49 | 35.24 | 2.45 | **0.014** | 2.49 | 9.61 | **< .001** |
| Vermis X | -2.35 | 3.67 | -0.64 | 0.522 | 2.90 | 5.72 | **< .001** |
| Bilateral Anterior Lobe | 874.04 | 100.42 | 8.70 | **< .001** | 4.04 | 11.92 | **< .001** |
| Left Anterior Lobe | 477.60 | 53.38 | 8.95 | **< .001** | 1.75 | 15.35 | **< .001** |
| Right Anterior Lobe | 405.71 | 54.02 | 7.51 | **< .001** | 4.72 | 13.42 | **< .001** |
| Bilateral Posterior Lobe | 732.50 | 564.83 | 1.30 | 0.195 | 2.31 | 127.36 | **< .001** |
| Left Posterior Lobe | 360.35 | 270.52 | 1.33 | 0.183 | 2.78 | 113.27 | **< .001** |
| Right Posterior Lobe | 276.12 | 279.40 | 0.99 | 0.323 | 2.30 | 134.80 | **< .001** |
| Bilateral Flocculonodular Lobe | 30.80 | 8.96 | 3.44 | **< .001** | 2.33 | 4.51 | **0.004** |
| Whole Vermis | 83.73 | 36.59 | 2.29 | **0.022** | 2.75 | 9.47 | **< .001** |
| Left Cerebellar Hemisphere | 822.79 | 297.09 | 2.77 | **0.006** | 2.89 | 102.37 | **< .001** |
| Right Cerebellar Hemisphere | 628.03 | 312.22 | 2.01 | 0.045 | 2.52 | 110.53 | **< .001** |
| Corpus Medullare | -592.36 | 93.87 | -6.31 | **< .001** | 3.43 | 100.46 | **< .001** |
| Total Cerebellum | 1283.73 | 648.17 | 1.98 | 0.048 | 3.17 | 124.94 | **< .001** |

*Note.* Sex (Male) = parametric fixed effect difference in volume for Males relative to Females (reference). s(age) = smoothed age effect on volume. Estimate represents the difference in volume (mm³) for Males compared to Females (reference group) after controlling for age, eTIV, and random scanner effects. Bold significance values indicate correction survival (p < 0.05). SE= Standard Error, edf = Effective Degrees of Freedom.

### Longitudinal Age Model

In the longitudinal model, after correction, 39 of the 41 regions displayed significant age effects. All the significant models illustrated that age is negatively associated with cerebellar subregional volume. The first derivative plots illustrate the heterogeneity of temporal variation of volume trends, with heightened atrophies commonly appearing between 40-60 years of life, demonstrated via **Fig. 2**. An important caveat is that since the main effect of age was modeled as age at each timepoint, rather than the common baseline age and time, the estimation reflects both between and within subject changes. Thus, when contextualizing the observed negative trend, it should not be classified as pure longitudinal volume rate change. See **Table 5** for further information per ROI parametric and smooth effects as well as **Supplemental Fig. 3** for derivative plots for all ROIs.

**Fig 2.**
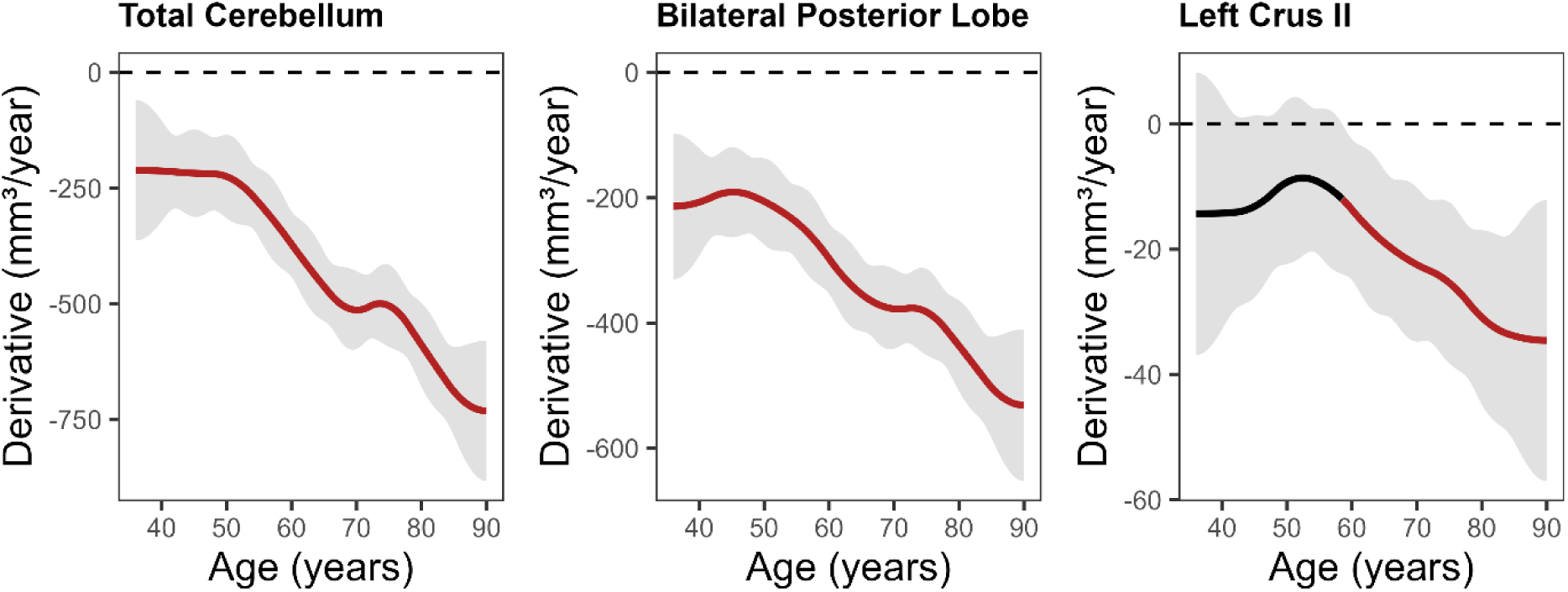
Panel of first derivative plots of smoothed age effects from the longitudinal models for Total Cerebellum, Bilateral Posterior Cerebellum, and Left Crus II volume, evaluating ages where the instantaneous rate of volume change (mm³/year) was significantly different from zero. Red segments indicate significance, where the 95% confidence interval grey band does not overlap with zero. The dashed horizontal line marks zero (no change in volume). The increased confidence intervals around age 90 may be attributed to the binning of participants who were reported to be 90 years of age or older.

**Table 5.** Sex and Age effects on cerebellar subregional volumes from the longitudinal model.

|  | ***Sex (Male)*** |  |  |  | ***s(age)*** |  |  |
|---|---|---|---|---|---|---|---|
| **Region** | **Estimate** | ***SE*** | ***t-value*** | ***p*** | ***edf*** | ***F*** | ***p*** |
| Left I-III | 18.92 | 10.29 | 1.84 | 0.066 | 2.69 | 3.59 | **0.012** |
| Left IV | 302.66 | 31.48 | 9.62 | **< .001** | 1.00 | 0.24 | 0.623 |
| Left V | 154.50 | 26.27 | 5.88 | **< .001** | 2.07 | 76.79 | **< .001** |
| Left VI | -331.51 | 82.13 | -4.04 | **< .001** | 5.26 | 73.17 | **< .001** |
| Left Crus I | 91.48 | 111.50 | 0.82 | 0.412 | 5.55 | 201.20 | **< .001** |
| Left Crus II | 229.79 | 79.15 | 2.90 | **0.004** | 3.64 | 40.38 | **< .001** |
| Left VIIB | -242.37 | 64.83 | -3.74 | **< .001** | 1.77 | 181.43 | **< .001** |
| Left VIIIA | 597.16 | 66.10 | 9.03 | **< .001** | 2.73 | 24.49 | **< .001** |
| Left VIIIB | -15.56 | 38.65 | -0.40 | 0.687 | 1.59 | 123.74 | **< .001** |
| Left IX | -39.03 | 40.29 | -0.97 | 0.333 | 3.25 | 62.29 | **< .001** |
| Left X | 20.19 | 4.67 | 4.32 | **< .001** | 2.86 | 3.84 | **0.010** |
| Right I-III | 11.32 | 10.43 | 1.09 | 0.278 | 2.03 | 17.45 | **< .001** |
| Right IV | 284.10 | 32.72 | 8.68 | **< .001** | 5.02 | 8.68 | **< .001** |
| Right V | 103.03 | 28.09 | 3.67 | **< .001** | 3.83 | 90.34 | **< .001** |
| Right VI | -332.67 | 76.20 | -4.37 | **< .001** | 4.64 | 67.97 | **< .001** |
| Right Crus I | 310.99 | 116.27 | 2.67 | **0.008** | 4.74 | 142.73 | **< .001** |
| Right Crus II | 305.03 | 80.71 | 3.78 | **< .001** | 1.65 | 59.86 | **< .001** |
| Right VIIB | -101.80 | 64.88 | -1.57 | 0.117 | 2.73 | 180.67 | **< .001** |
| Right VIIIA | 172.64 | 61.13 | 2.82 | **0.005** | 2.17 | 1.98 | 0.111 |
| Right VIIIB | -26.20 | 36.56 | -0.72 | 0.474 | 3.52 | 104.23 | **< .001** |
| Right IX | 14.33 | 39.95 | 0.36 | 0.720 | 2.35 | 84.97 | **< .001** |
| Right X | 11.03 | 4.87 | 2.27 | **0.024** | 2.14 | 13.41 | **< .001** |
| Vermis VI | 12.15 | 13.38 | 0.91 | 0.364 | 3.12 | 30.21 | **< .001** |
| Vermis VII | -0.90 | 10.24 | -0.09 | 0.930 | 3.75 | 15.54 | **< .001** |
| Vermis VI-VII | 9.90 | 19.31 | 0.51 | 0.608 | 3.61 | 36.50 | **< .001** |
| Vermis VIII | 115.74 | 19.31 | 5.99 | **< .001** | 2.69 | 7.56 | **< .001** |
| Vermis IX | -39.54 | 9.10 | -4.34 | **< .001** | 3.78 | 24.75 | **< .001** |
| Vermis VI-IX | 79.15 | 34.60 | 2.29 | **0.022** | 3.73 | 42.88 | **< .001** |
| Vermis X | -3.21 | 3.55 | -0.90 | 0.367 | 3.65 | 10.96 | **< .001** |
| Bilateral Anterior Lobe | 816.73 | 97.93 | 8.34 | **< .001** | 4.54 | 38.79 | **< .001** |
| Left Anterior Lobe | 456.25 | 51.76 | 8.81 | **< .001** | 2.86 | 23.59 | **< .001** |
| Right Anterior Lobe | 374.14 | 53.02 | 7.06 | **< .001** | 4.99 | 43.79 | **< .001** |
| Bilateral Posterior Lobe | 615.16 | 541.86 | 1.14 | 0.256 | 6.07 | 280.19 | **< .001** |
| Left Posterior Lobe | 143.83 | 265.57 | 0.54 | 0.588 | 6.37 | 275.78 | **< .001** |
| Right Posterior Lobe | 304.02 | 268.23 | 1.13 | 0.257 | 5.77 | 248.87 | **< .001** |
| Bilateral Flocculonodular Lobe | 31.65 | 8.79 | 3.60 | **< .001** | 2.71 | 8.62 | **< .001** |
| Whole Vermis | 75.46 | 35.91 | 2.10 | 0.036 | 3.90 | 45.17 | **< .001** |
| Left Cerebellar Hemisphere | 631.36 | 288.44 | 2.19 | 0.029 | 6.23 | 307.27 | **< .001** |
| Right Cerebellar Hemisphere | 645.17 | 297.16 | 2.17 | 0.030 | 5.91 | 263.21 | **< .001** |
| Corpus Medullare | -539.26 | 92.34 | -5.84 | **< .001** | 5.29 | 120.91 | **< .001** |
| Total Cerebellum | 1220.77 | 628.80 | 1.94 | 0.052 | 6.64 | 316.98 | **< .001** |

*Note.* Sex (Male) = parametric fixed effect difference in volume for Males relative to Females (reference). s(age) = smoothed age effect on volume. Estimate represents the difference in volume (mm³) for Males compared to Females (reference group) after controlling for age, eTIV, and random scanner effects. Bold significance values indicate correction survival (p < 0.05). SE= Standard Error, edf = Effective Degrees of Freedom.

### Sex*Age Interaction Model

The interaction of sex and age was significant in 17 of the 41 regions after correction. All significant regions exhibited linear deviations except for Left Crus I, which suggested a quadratic deviation. Regions that were insignificant suggest that while there may be differences in volume level, see **Table 5**, the trajectories across age between both sexes are similar. Contrarily, regions that exhibited significant linear deviations suggest that there is a constant divergence in volume, where one sex possess greater volumetric reductions across age. For Left Crus I, the quadratic-like deviation indicates that there is a distinct age pattern between sexes, rather than just differences in rate of change. Additional information of the nonparametric sex and age effects is observable in **Table 6**. Examination of **Fig. 3** showcases the deviation shape for Left Crus I compared to Total Cerebellum to visualize linear and nonlinear effects (See **Supplemental Fig. 4** for all ROI partial effect plots and estimated trend plots parsed by sex).

**Fig 3.**
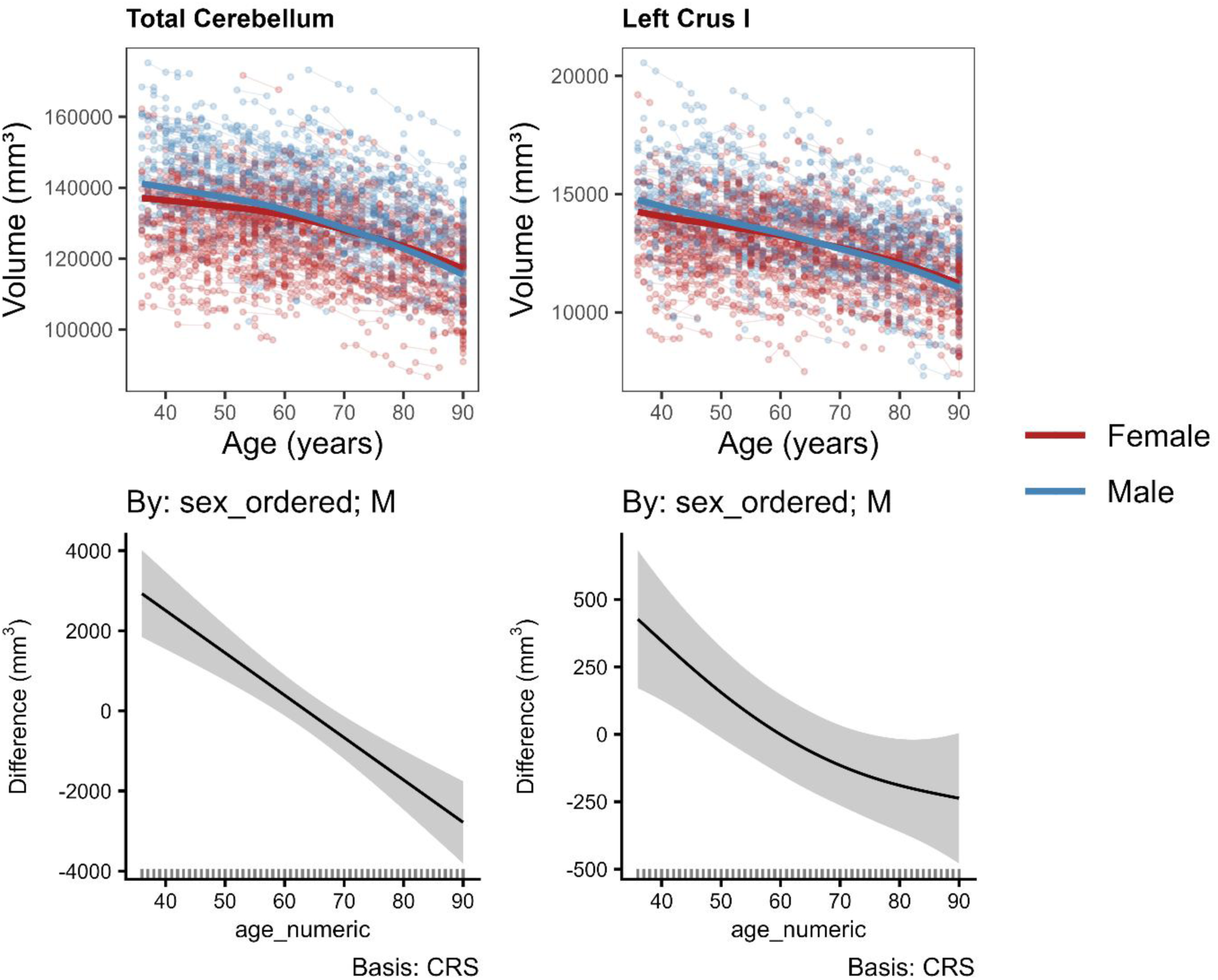
Top row shows predicted model trajectories for males (blue) and females (red) from the sex*age interaction model overlaid on uncorrected volumetric data across visits for Total Cerebellum and Left Crus I. Bottom row show partial effect plots illustrating how volumetric trends (mm³) may deviate for males relative to the mean from females (reference) across age. Values at zero indicate no difference; positive values are indicative of greater male volume at the respective age, and vice versa for negative values. Each partial effect plot was derived from the sex*age interaction model, paired with 95% confidence interval grey bands. Smooth interaction terms used cubic regression splines as the basis function (Basis: CRS). Linear deviations illustrate a linear deviation from reference (i.e., edf = 1.00) while nonlinear effects indicate nonlinear deviations (i.e., edf = 2.00). Models controlled for eTIV, and random subject and scanner effects.

**Table 6.** Interaction Between Sex and Age Effects on Cerebellar Subregional Volumes from the Sex*Age Interaction Model.

|  | ***s(age) × sex*** |  |  |
|---|---|---|---|
| **Region** | ***edf*** | ***F*** | ***p*** |
| Left I-III | 2.67 | 3.46 | 0.063 |
| Left IV | 1.00 | 7.07 | **0.008** |
| Left V | 1.00 | 3.04 | 0.081 |
| Left VI | 1.29 | 16.78 | **< .001** |
| Left Crus I | 2.04 | 7.47 | **< .001** |
| Left Crus II | 1.00 | 3.27 | 0.071 |
| Left VIIB | 1.00 | 0.87 | 0.350 |
| Left VIIIA | 1.88 | 0.94 | 0.394 |
| Left VIIIB | 1.00 | 2.51 | 0.113 |
| Left IX | 1.00 | 0.00 | 0.953 |
| Left X | 1.78 | 1.38 | 0.361 |
| Right I-III | 1.00 | 0.14 | 0.704 |
| Right IV | 1.00 | 6.57 | **0.010** |
| Right V | 1.38 | 1.46 | 0.155 |
| Right VI | 1.00 | 18.21 | **< .001** |
| Right Crus I | 1.00 | 14.90 | **< .001** |
| Right Crus II | 1.77 | 4.53 | 0.043 |
| Right VIIB | 1.00 | 3.89 | 0.049 |
| Right VIIIA | 1.00 | 0.90 | 0.344 |
| Right VIIIB | 2.60 | 1.37 | 0.153 |
| Right IX | 1.00 | 0.64 | 0.424 |
| Right X | 1.00 | 2.54 | 0.111 |
| Vermis VI | 1.00 | 2.30 | 0.129 |
| Vermis VII | 1.00 | 2.10 | 0.147 |
| Vermis VI-VII | 1.00 | 4.21 | 0.040 |
| Vermis VIII | 1.00 | 3.42 | 0.064 |
| Vermis IX | 1.00 | 3.05 | 0.081 |
| Vermis VI-IX | 1.00 | 8.27 | **0.004** |
| Vermis X | 1.00 | 1.13 | 0.288 |
| Bilateral Anterior Lobe | 1.00 | 5.07 | 0.024 |
| Left Anterior Lobe | 1.00 | 6.18 | **0.013** |
| Right Anterior Lobe | 1.00 | 5.37 | **0.021** |
| Bilateral Posterior Lobe | 1.00 | 36.09 | **< .001** |
| Left Posterior Lobe | 1.00 | 24.78 | **< .001** |
| Right Posterior Lobe | 1.00 | 28.04 | **< .001** |
| Bilateral Flocculonodular Lobe | 1.00 | 1.58 | 0.209 |
| Whole Vermis | 1.00 | 8.17 | **0.004** |
| Left Cerebellar Hemisphere | 1.00 | 30.90 | **< .001** |
| Right Cerebellar Hemisphere | 1.00 | 32.30 | **< .001** |
| Corpus Medullare | 1.00 | 15.25 | **< .001** |
| Total Cerebellum | 1.00 | 35.69 | **< .001** |

*Note.* The factor interaction of s(age) × sex tests whether sex alters aging trajectory. Each model controlled for age, eTIV, and random subject and scanner effects. Bold p values indicate FDR correction survival (p < 0.05). edf = Effective Degrees of Freedom.

### Sensitivity Models

Sensitivity models identical to the primary models were run, with each confounder added independently to assess their impact on the main effects. Summaries of analytic outcomes can be found in **Table 7**. Additionally, comparative metrics between primary and sensitivity models are documented in **Supplemental Table 2**.

**Table 7.** Summary of ROIs that had altered significance after sensitivity analyses across models.

| **Model** | **Gained Significance** | **Lost Significance** |
|---|---|---|
| *Cross-sectional Model* |  |  |
| Blood Pressure Classification | - | Left I-III |
| Race | Vermis VIII | Left I-III, Vermis VII |
| Ethnicity | Left X | - |
| Education | - | Left I-III |
| MoCA | - | Bilateral Flocculonodular Lobe |
| *Longitudinal Age Model* |  |  |
| Blood Pressure Classification | - | - |
| Race | Right VIIIA | - |
| Ethnicity | - | - |
| Education | - | Vermis VIII |
| MoCA | - | Left X |
| *Sex\*Age Interaction Model* |  |  |
| Blood Pressure Classification | - | - |
| Race | Left I-III, Left Crus II | Vermis VI-VII, Right VIIB |
| Ethnicity | - | Vermis VI-VII, Right VIIB |
| Education | - | Right Crus II, Right VIIB |
| MoCA | Vermis VIII | Right Crus II, Right VIIB |

*Note.* If a ROI is not listed, no change in significance occurred after the sensitivity analysis. Significance is defined as *p* < 0.05.

## Discussion

In this work we modeled nonlinear cerebellar subregional volumetric trends across healthy aging adults. Cross-sectional results indicate that older adults present reduced volume for 34 of the 41 ROIs compared to younger adults, with the anterior cerebellum being less affected by age. Longitudinal analyses revealed significant negative associations with age for 39 of the 41 regions, with heightened age effects becoming evident around midlife and predominantly in the neocerebellum. For significant sex differences, men consistently displayed overall larger volumes compared to women across the cerebellum in the cross-sectional and longitudinal models. However, select significant regions from the interaction model showed male trajectories crossed below those of females at older ages, see **Supplemental Fig. 4**. Furthermore, when longitudinally modeling the interaction between sex and age, 16 of the segmentations featured linear volume trajectory deviations, while Left Crus I had a quadratic deviation between men and women. Taken together, cerebellar senescence is suggested to be a heterogeneous phenomenon that appears to robustly affect posterior regions.

Across models, negative age effects were observed. A greater proportion of significant age effects were consistently seen in posterior neocerebellar regions-many of which were nonlinear. This is suggestive of regional aging susceptibility and provides evidence for volumetric heterogeneity. In agreement with previous work (d’Oleire Uquillas et al., 2026; Han et al., 2020a; Romero et al., 2021; Wang et al., 2024), these findings may help explain age-related cognitive decline as well illustrate why they consistently appear affected in various neuropathologies (Chen et al., 2019; Chen et al., 2024; Rashidi et al., 2026). Additionally, while overall negative bilateral patterns were prominent, many displayed unique lateralized trends, adhering to the notion of cerebellar regional specificity (Cerminara et al., 2015; d’Oleire Uquillas et al., 2026). Interestingly, no age effects were observed for Left IV across models, suggesting resistance to aging. In Right IV, while eventually there were degenerative trends later in life, significant increases in volume were observed in midlife, see **Supplemental Fig. 3 and Supplemental Table 1**. This contrasts with cross-sectional (d’Oleire Uquillas et al., 2026; Wang et al., 2024) and lifespan (Romero et al., 2021) work where negative bilateral age effects for Lobule IV were seen. However, core modeling approach deviations exist which prevent direct comparison of the present results. Even so, similar longitudinal work from Han et al. (2020a) observed positive bilateral volumetric trends in Lobule IV across time. While not in total agreement, both findings allude to Lobule IV possessing a distinct age-related morphologic trajectory. Taken together, the overall results showed an increased detection of age effects across regions and as well as pronounced first derivative magnitudes via the longitudinal models.

Consistent with the notion that males typically have larger brains and grey matter (Ritchie et al., 2018), the majority of our results adhere to this trend. This deviates with recent work from Hicks et al. (2022) where most of their significant findings suggested women have higher regional volumes. However, it is worth noting that while several studies have explored cerebellar sex differences, across lobules the results remain inconsistent (Han et al., 2020a, Hicks et al. 2022; Luft et al., 1999; Romero et al., 2021; Steele and Chakravarty et al. 2018; Wang et al., 2024; Weier et al. 2014). A possible explanation is that many studies report sex differences based on either absolute cerebellar volume with no TIV control or model cerebellar volumes as a proportion of total head size, assuming a 1:1 ratio and confounding sex differences. Similar to previous longitudinal work of Han et al. (2020a), we introduced TIV as a covariate rather than computing proportional volumes, avoiding potential confounding sex differences in head-size differences. With that said, robust findings were found in agreement with several previous studies in which men possessed greater bilateral Lobule VIIIA volume (Hicks et al., 2022; Steele and Chakravarty et al. 2018; Wang et al., 2024) and women had greater bilateral Lobule VI volume (Hicks et al., 2022). In a more appropriate comparison, while Han et al. (2020a) reported sparse sex differences, they also found men exhibited larger volumes for Lobule VIIIA and Vermis IX. Moreover, when assessing volumetric deviation trends by sex, they found Right Crus II and Corpus Medullare have steeper atrophy patterns in men while women displayed steeper negative trajectories in the Left Anterior Lobe (Han et al., 2020a); all but Right Crus II are in agreement with our findings. Overall, the inconsistency of findings paired with exploratory evidence of sex differences in patterns of cerebellar decline, warrant further research with consistent modeling methods across independent datasets.

The reason for the greater prevalence of age effects on the posterior cerebellar structures may be multifaceted. Limited work exists that investigate the present phenomenon, and the subsequent discussion of potential conceptual frameworks is speculative. One of the few studies that commented on the posterior aging bias (d’Oleire Uquillas et al., 2026) attributed their related findings to the retrogenesis hypothesis (Reisberg et al., 1999). Retrogenesis, typically applied to neurodegenerative disorders like Alzheimer’s, posits that higher cognitive order nodes are more prone to degeneration than are resilient motor related nodes-which align with the observed results, as the posterior cerebellum is seen as the cognitive center. This is complimented by Guo et al. (2016), where they introduced the cerebro-cerebellar connectivity-based degeneration hypothesis, suggesting that cerebellar atrophy adheres to preferential degeneration of regions that are involved in altered cerebral networks commonly associated with neurodegenerative diseases such as the default mode and salience networks. Cellular and preclinical work appears to be in disagreement with these findings, suggesting that anterior regions are at heightened risk of neurodegeneration, attributed to the presence of higher Zebrin II negative zones per capita than posterior regions (Cerminara et al., 2015; Lin et al., 2020; Zhou et al. 2014). These studies however are not directly comparable, as they specifically examine Purkinje cell loss, rather than gross subregional morphometry. Moreover, Zebrin II zones do not perfectly align with commonly used ROIs in neuroimaging research.

To bridge this gap between these contrasting ideas, further mechanistic studies are necessary to disentangle potential biological drivers behind the age vulnerability of posterior lobules. One such avenue that may aid in the elucidation of the present findings are investigations of potential regional neuroinflammatory effects, given that cerebellar microglia are prone to immune-vigilant states (Grabert et al., 2016), which may prove to be aversive as chronic inflammation is a hallmark of aging (Li et al., 2023; Singh et al., 2025). Relatedly, there is increased recognition that cellular senescence is a biological hallmark of aging. There has been recent postmortem tissue work showing spatial heterogeneity of senescent signatures across cell-types and layers of the dorsal lateral prefrontal cortex (Sloan et al., 2026). The significance of this is that future work could apply similar techniques to the cerebellum to characterize whether different senescent profiles across cerebellar regions exist and if such differences may explain regional patterns of age-related atrophy. Taken together, neocerebellar regions seem to be at an increased risk for age effects, but further investigations into their spatially unique degenerative etiology is needed.

There is growing consensus that middle-age is a critical phasic transitionary period in life. Dohm-Hansen and colleagues (2024) argue that studies who encapsulate broad age ranges should employ nonlinear modeling to capture a myriad of nuanced biological processes. In the present study, pronounced nonlinear transitions in degenerative pathways across the cerebellum were observed, particularly during midlife. Similar dynamic trends have been found in canonical age-susceptible regions (Luo et al., 2019; Schuff et al., 2012). For example, Luo et al. (2019) reported that nonlinear trajectory shifts in grey matter volume trends for the parahippocampus appeared around 40 years of age. Additionally, the demonstrated plateauing effect in late-life, thought as a potential age aggregation artifact, also has been reported in cerebral regions (Schuff et al., 2012), and relatedly for absolute volumes of bilateral Crus II and Lobule X as well (Romero et al., 2021). Moreover, cerebellar lifespan work from Romero et al. (2021) further illustrated the criticality of midlife, where various transitionary periods to volumetric reductions started for posterior regions. These results are clinically complemented by Zhou et al. (2023), where they found dual gait task performance began to decline in late midlife healthy adult-aligning with the historical role of the cerebellum in motor coordination, gait, and ataxia-related disorders (Harding et al., 2021). Moreover, the pronounced degenerative rate in middle age may shed light on other findings that found marked decline in cognitive and behavioral performance in middle age (Statsenko et al., 2021).

As previously mentioned, GAMMs are a modeling technique derived from traditional linear approaches that can examine nonlinear trends. Future work investigating both cross-sectional and longitudinal morphometry would benefit from this approach to prevent a linear constraint of biological trends that otherwise may be dynamic in nature. Moreover, the present study benefited from fine cerebellar grey matter segmentation via ACAPULCO (Han et al., 2020b; Kerestes et al. 2022), to investigate subregional aging morphometry, compared to traditional cerebellar approaches. Beyond this, the present study employed a longitudinal design allowing for the examination of between subject and within-subject nonlinear changes, providing greater insight into the proposed variable age-related structural patterns. Thereby leveraging analogous techniques for aging populations may prove to be beneficial for the cerebellum and cerebrum alike.

In summary, these results suggest that the cerebellum is markedly affected by age, in a heterogenic pattern where region specific volumetric reduction patterns exist. The relevancy of these findings is underscored by the expansion of the aging population as well as the increased recognition of the role of the cerebellum in neurodegenerative disorders. Furthermore, the pronounced atrophy detected in midlife may aid in intervention strategies to know when opportune times to stave off age-related cognitive decline exist. Another important takeaway is the conceptualization of the areas that were identified at risk of age-related degeneration. Here, regions commonly associated with cognitive processes were consistently associated with negative age effects, highlighting the significance of these findings. Overall, results from the present investigation may provide a foundation for future investigations examining cerebellar alterations across both neurodegenerative and neuropsychiatric disorders.

## Limitations

Several limitations exist in the present study that should be of note. First, ACAPULCO is not adept to fully assess true cerebellar white matter, leading to potential under inclusion. Additionally, the segmentation does not include deep cerebellar nuclei, which are the output hubs of the cerebellum. Secondly, while GAMMs enable modeling smooth functions for nonlinear trends, this can lead to potential overfitting. However, measures were taken to reduce the risk of overfitting. Such measures included employing penalized smooth terms via REML estimation, assessing quantile-quantile plots of model residuals, conducting basis dimension diagnostics, and running sensitivity analyses. Building off this, smooth terms may be difficult to intercept, thereby leading to the need for inferential estimates to examine rates of change and directionality. Lastly, like previous longitudinal work (Han et al., 2020a), our sample was overly Caucasian. As a result, this may limit the generalizability of the present findings in dementia-related research, where demographic groups such as Hispanic populations face a rising prevalence of dementia (Llibre-Guerra et al., 2026). Future work should investigate deep cerebellar nuclei morphometry trends as well as continue nonlinear modeling efforts considering various disorder profiles and representative samples.

## Conclusion

In the present work, we extend conventional knowledge on cerebellar morphometry across aging by investigating the nonlinear trends in a region-specific manner. Overall, we found a variable negative association of age spanning the cerebellum, with marked midlife grey matter reduction. Through the continued investigation of age-related structural and functional trends of the cerebellum, foundational insight into the cerebellar-cognitive interplay can be achieved.

## Acknowledgements

Data reported in this publication were provided in whole by the Aging Adult Vulnerability and Resiliency in the Aging Adult Brain Connectome (AABC) project (U19AG073585) and the Human Connectome Project in Aging (HCP-A, U01AG052564) funded by the National Institute of Aging of the National Institutes of Health. HCP-A was further supported by funds provided by the McDonnell Center for Neuroscience at Washington University in St. Louis.

The HCP-Aging and AABC Release 2.0 data used in this report came from ConnectomeDB powered by BALSA: https://balsa.wustl.edu/project?project=AABC

## Author Contributions

Z.M.B led the conceptualization of the project, conducted data processing, data quality assurance, data analysis, and wrote the manuscript. N.D contributed to data quality assurance and the literature review of the manuscript. J.A.T. provided funding to support this research and supervision. All authors reviewed and approved the manuscript.

## Declarations

### Ethics approval

Data for this study was obtained from the Human Connectome Project in Aging (HCP-A) and a longitudinal extension, Aging Adult Brain Connectome (AABC) Study Release 2.0 (grant numbers U01AG052564, U19AG073585). At each site, subjects underwent consent procedures as well as assessment for consent capacity. No new data was collected for the present study. No additional institutional review board (IRB) approval was required per The Ohio State University’s regulatory policies.

