## Supplement for "Longitudinal examination of cerebellar morphometry across typical aging adults reveals nonlinear heterogenous trends"

***Supplemental Information***

**Supplemental Methods**

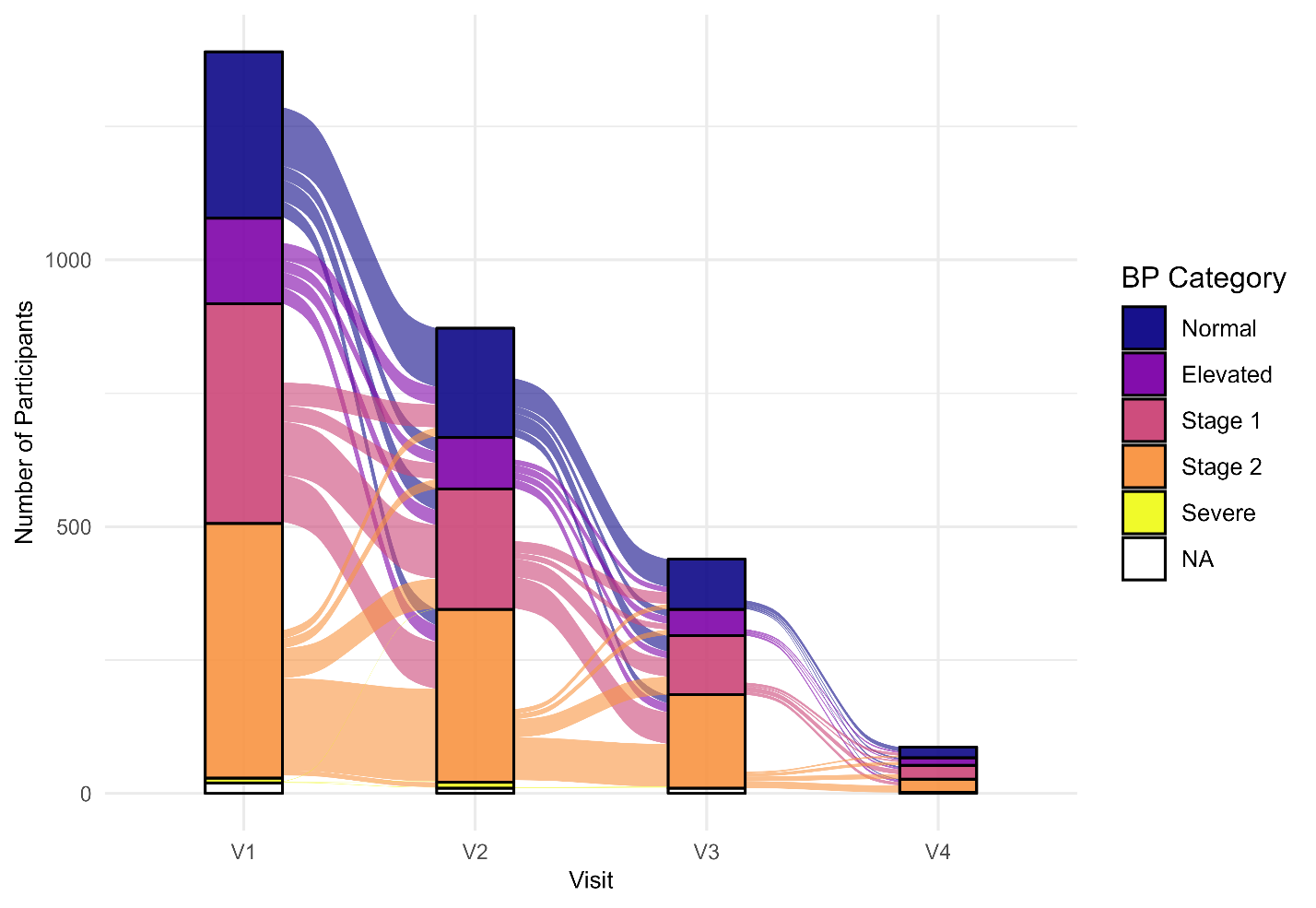

**Supplemental Fig. 1.** Plot showing blood pressure classification across visits illustrating relative consistency across visits. Color band width for within and between visit distributions represents the proportion of participants within a given blood pressure classification. Category designation previously defined in the associated manuscript.

*First derivative calculation*

The purpose of generating first derivative plots for the fitted smooth age effects was to examine nonlinear volumetric changes for each region across the observed age span. To accomplish this, the package *gratia* (v0.11.2) was used. Moreover, this procedure was done only for cross-sectional and longitudinal age models. In order to explore the rate of change of the isolated age effect from the fitted models, the first derivatives were evaluated at 1,000 evenly spaced points across the age range (0.054 years/point) via forward finite differences. Reasoning behind this methodological choice was to ensure adequate plotting resolution for derivative trend visualization as well as precise identification of significant age windows. Equally as important, because the longitudinal age models used the *gamm4* package (v0.2.7), there is no calculated uncertainty matrix for the spline wiggliness. The importance of this is that there is an inability for the later discussed 95% confidence intervals (CIs) to account for selection-corrected Bayesian covariance matrix to account for the uncertainty involved in estimating the spline's penalty parameter. Therefore, both cross-sectional and longitudinal models did not use this function for comparability. When plotting the computed first derivatives (See **Supplemental Fig. 2** and **Fig. 3**), simultaneous 95% confidence intervals (CI) were calculated to examine which ages the associated derivative was significantly non-zero, in essence, indicating whether a significant increase or decrease in volume (mm³ per year) at that age exists. An insignificant result, where the CI passes zero indicates there is no significant age-related change in volume at that specific age. Of note, significant flat trends indicate a constant rate of change across each year- corroborated by their fitted model effective degrees of freedom (edf) value. In the subsequent first derivate plots, the x-axis denotes age while the y-axis indicates the estimated first derivative (mm³ per year), reflecting the slope of the fitted trajectory at any of the 1,000 evaluated points.

**Supplemental Results**

*Model assumptions check*

Each model underwent normality checking via visual inspection of residual histograms and Q-Q plots. No violations of normality were detected, and we concluded that the residuals displayed sufficient normal distribution. Moreover, assessment of the adequacy of the chosen basis function (k) were conducted for each model.

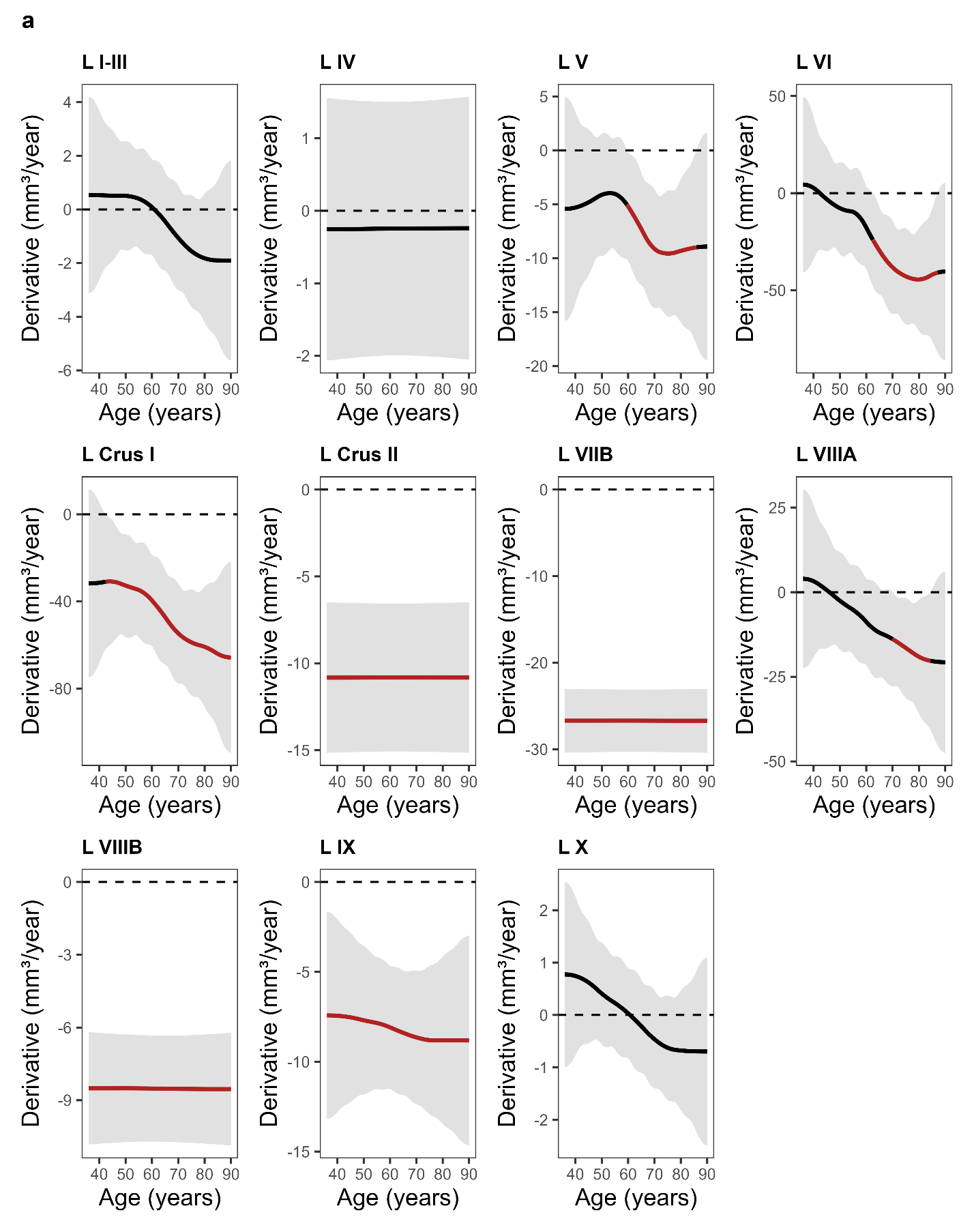

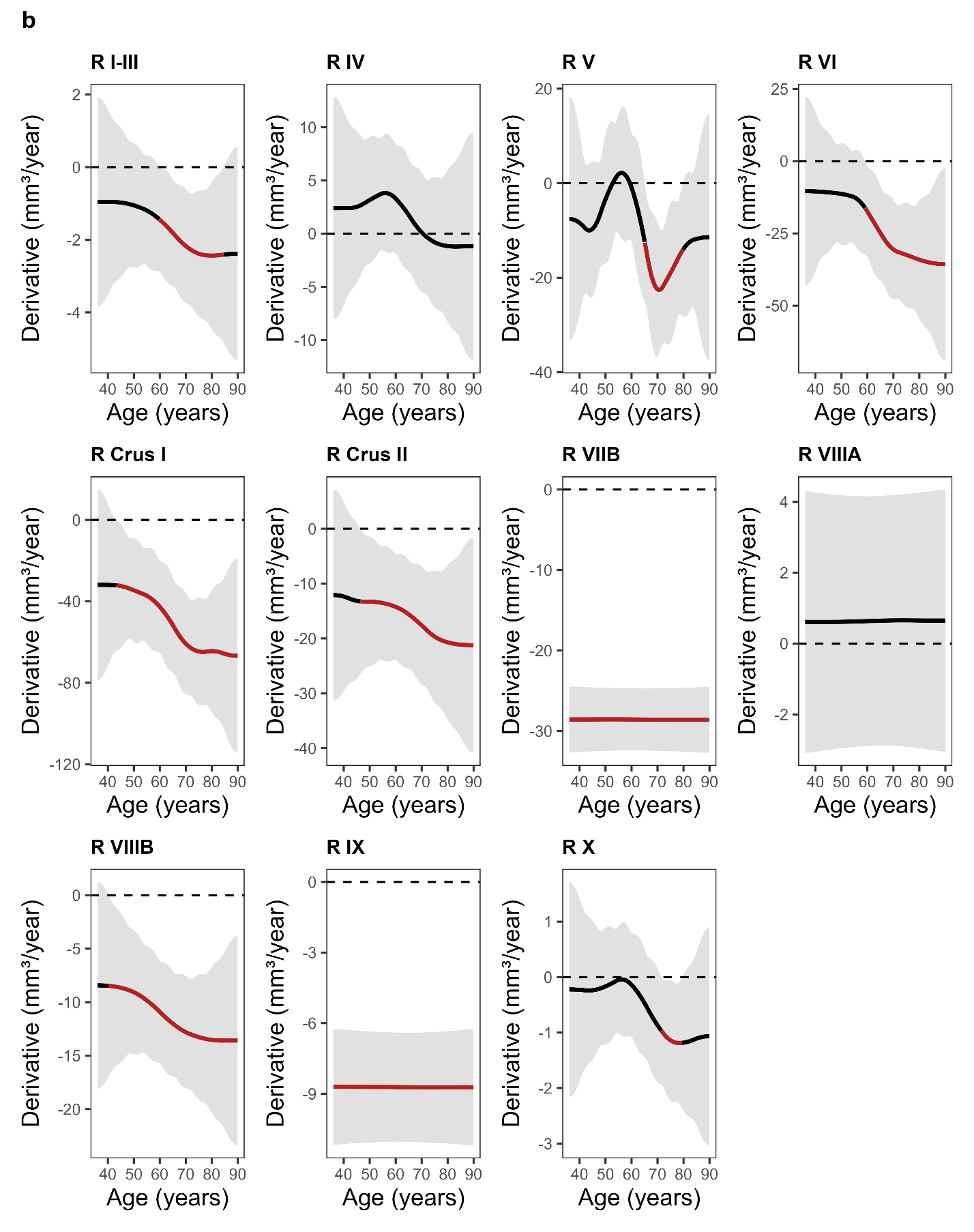

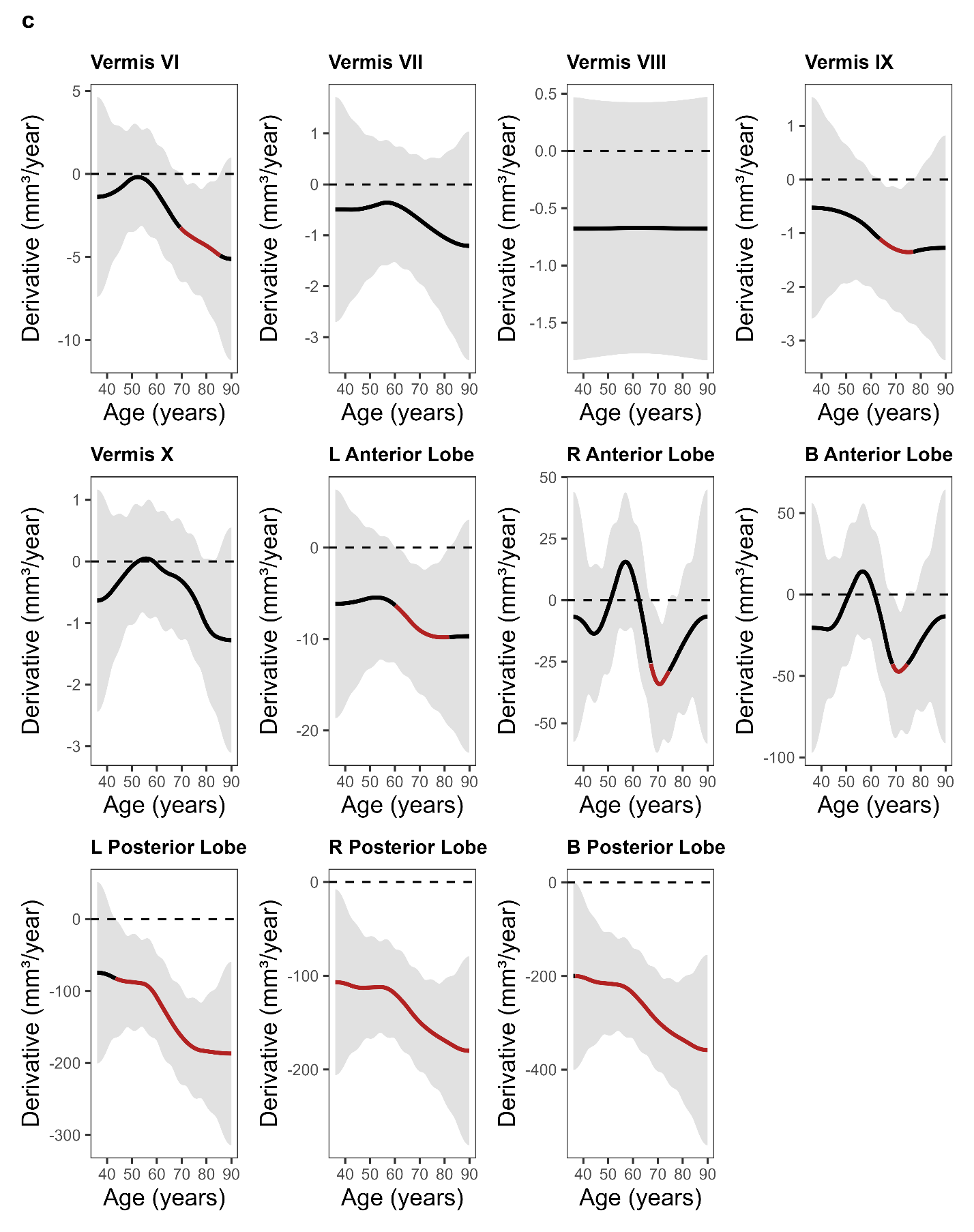

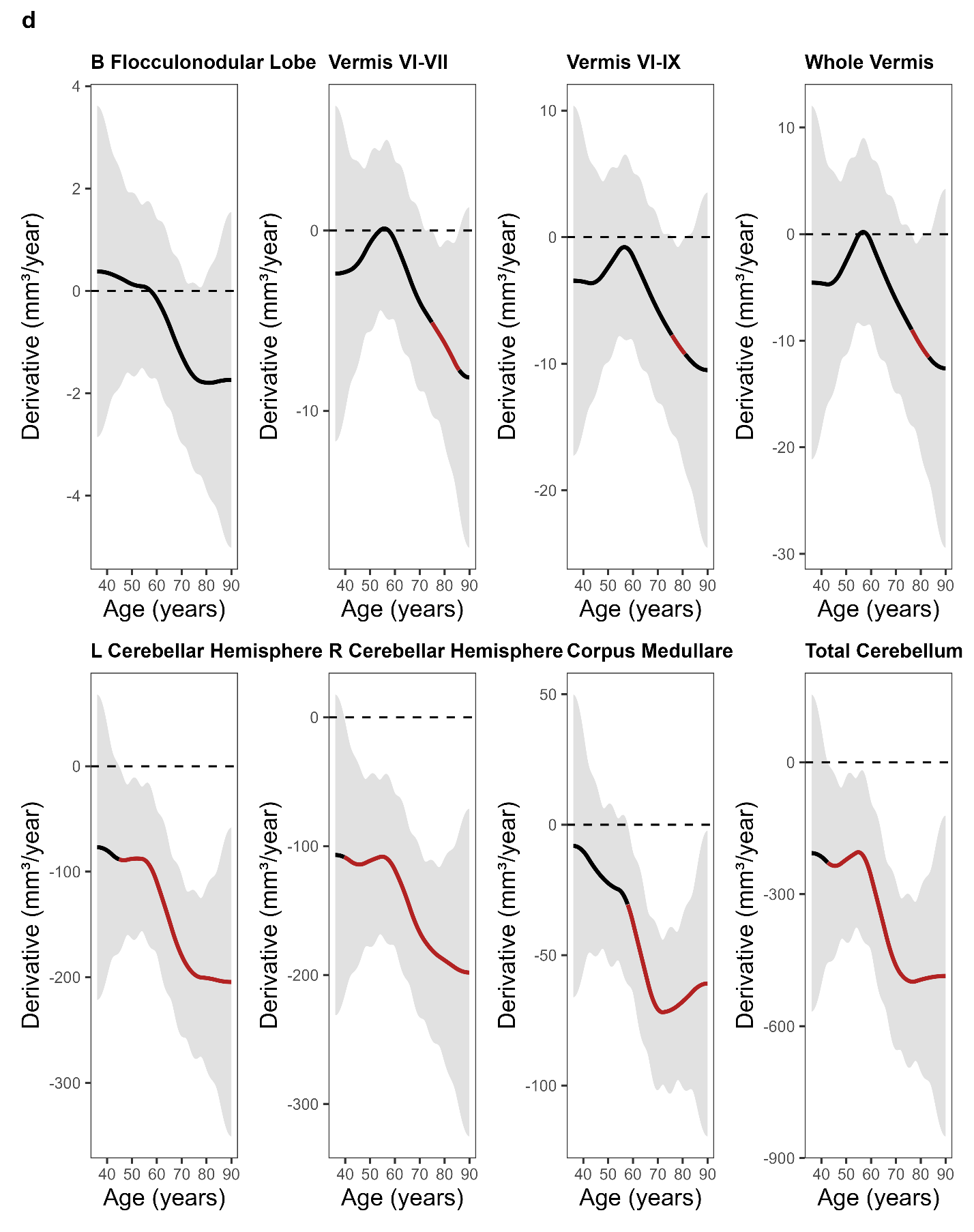

**Supplemental Fig. 2**. Grids of first derivative plots of smoothed age effects from the cross-sectional models evaluating ages where the instantaneous rate of volume change (mm³/year) was significantly different from zero. Red segments indicate significance, where the 95% confidence interval grey band does not overlap with zero. The dashed horizontal line marks zero (no change in volume). The plateauing effect around age 90 may be attributed to the binning of participants who were reported to be 90 years of age or older. ROIs were split into 4 grids (**Supplemental** **Fig. 2a** – **Supplemental** **Fig. 2d**) for easier visualization. Abbreviations in the plot titles are as follows: R = Right, L = Left, B = Bilateral.

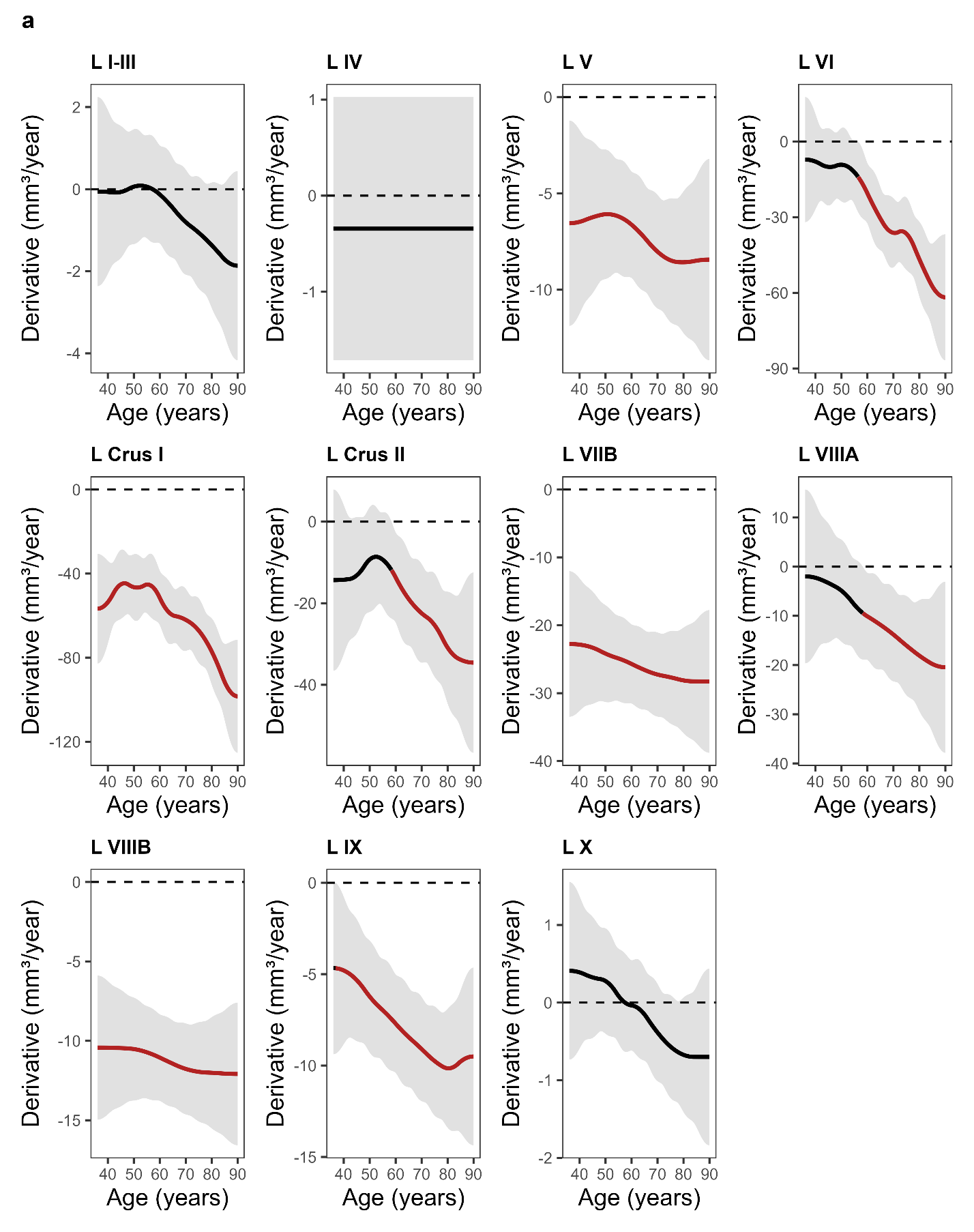

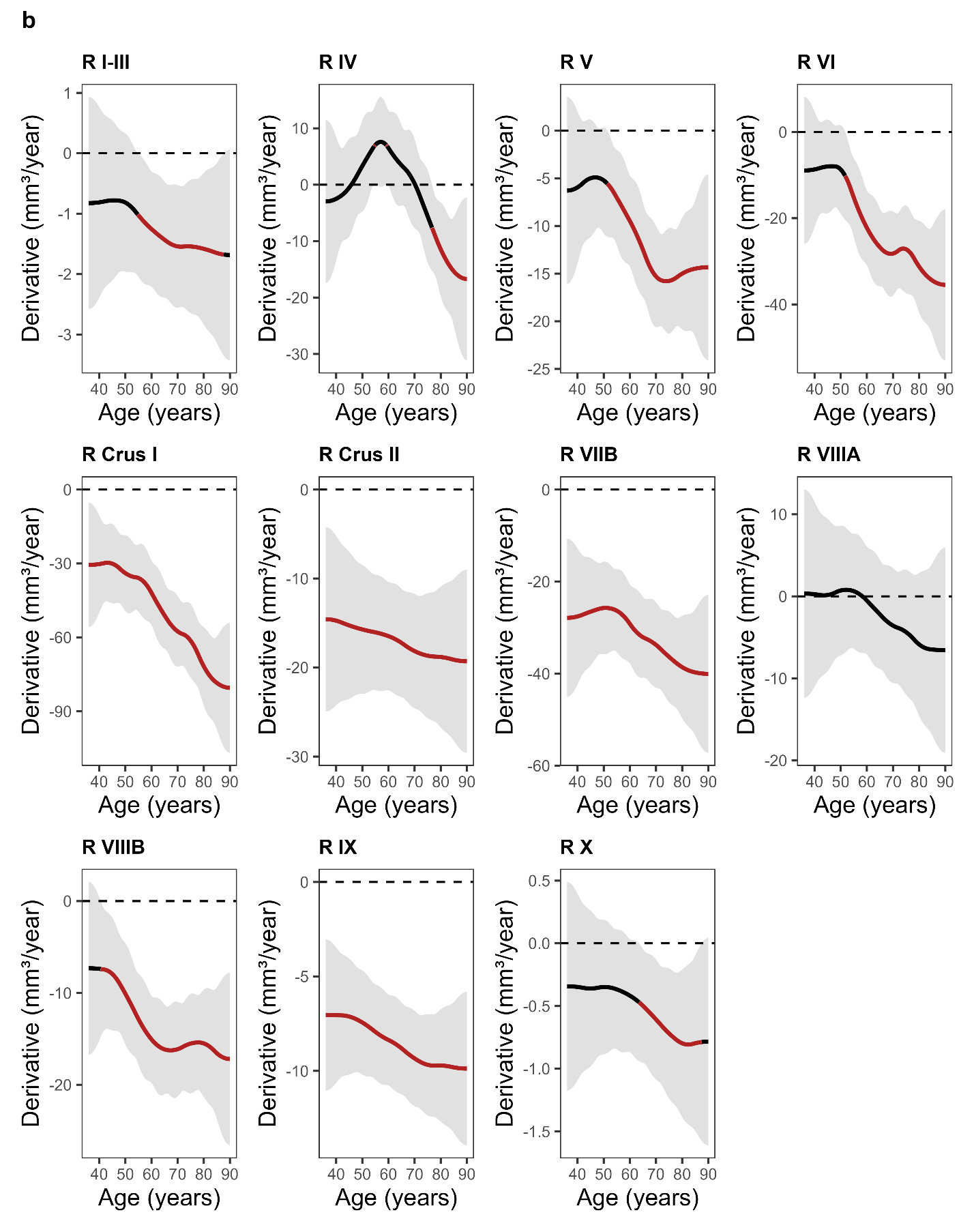

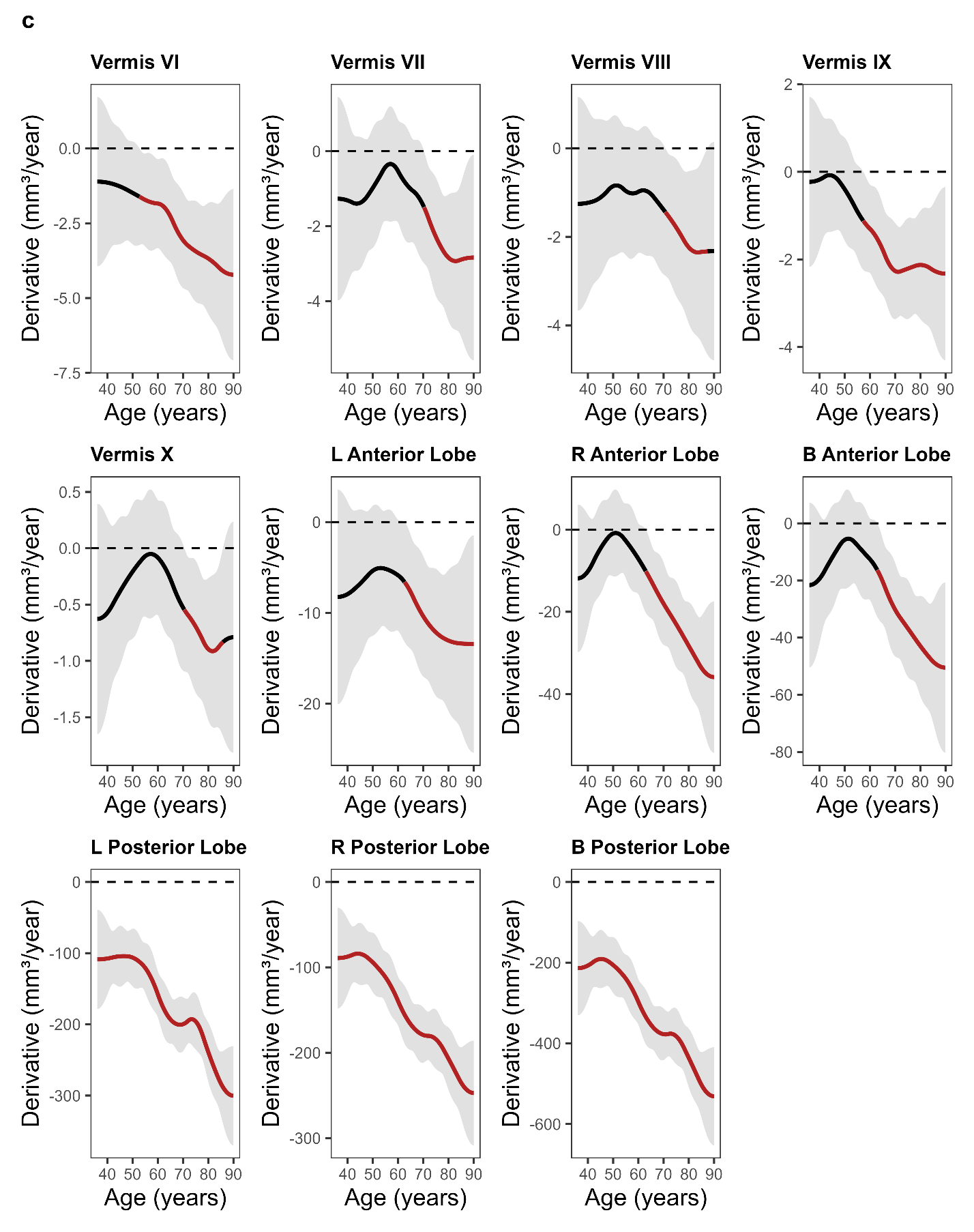

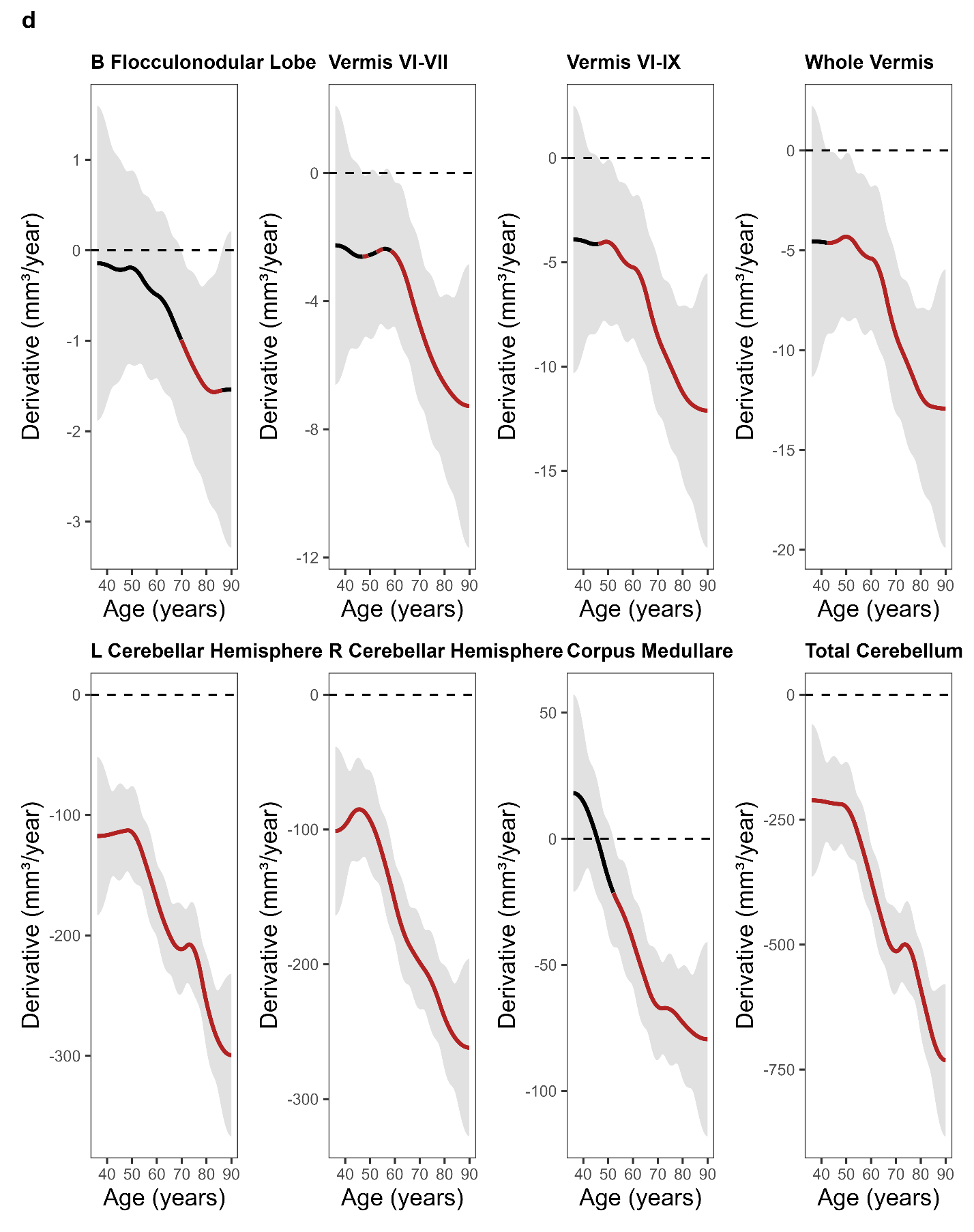

**Supplemental Fig. 3.** Grids of first derivative plots of smoothed age effects from the longitudinal models evaluating ages where the instantaneous rate of volume change (mm³/year) was significantly different from zero. Red segments indicate significance, where the 95% confidence interval grey band does not overlap with zero. The dashed horizontal line marks zero (no change in volume). The plateauing effect around age 90 may be attributed to the binning of participants who were reported to be 90 years of age or older. ROIs were split into 4 grids (**Supplemental** **Fig. 3a** – **Supplemental** **Fig. 3d**) for easier visualization. Abbreviations in the plot titles are as follows: R = Right, L = Left, B = Bilateral.

**Supplemental Table 1.** Ages where significant smoothed age effect changes occurred.

| **Region** | **Significant Age Range (years)** | **Peak Rate (mm³/year)** | **Direction** |
| --- | --- | --- | --- |
| Left I-III | n.s. | — | — |
| Left IV | n.s. | — | — |
| Left V | 36.0–90.0 | -8.57 | Decrease |
| Left VI | 56.3–90.0 | -61.75 | Decrease |
| Left Crus I | 36.0–90.0 | -98.39 | Decrease |
| Left Crus II | 58.5–90.0 | -34.56 | Decrease |
| Left VIIB | 36.0–90.0 | -28.28 | Decrease |
| Left VIIIA | 58.3–90.0 | -20.45 | Decrease |
| Left VIIIB | 36.0–90.0 | -12.09 | Decrease |
| Left IX | 36.7–90.0 | -10.15 | Decrease |
| Left X | n.s. | — | — |
| Right I-III | 54.9–87.6 | -1.68 | Decrease |
| Right IV | 54.5–55.1; 59.0–59.6; 76.6–90.0 | 7.11; 7.12; -16.70 | Increase; Decrease |
| Right V | 51.4–90.0 | -15.79 | Decrease |
| Right VI | 51.7–90.0 | -35.46 | Decrease |
| Right Crus I | 36.0–90.0 | -80.47 | Decrease |
| Right Crus II | 36.0–90.0 | -19.29 | Decrease |
| Right VIIB | 36.0–90.0 | -40.05 | Decrease |
| Right VIIIA | n.s. | — | — |
| Right VIIIB | 40.6–90.0 | -17.19 | Decrease |
| Right IX | 36.0–90.0 | -9.88 | Decrease |
| Right X | 63.7–87.2 | -0.81 | Decrease |
| Vermis VI | 52.8–90.0 | -4.22 | Decrease |
| Vermis VII | 70.3–90.0 | -2.93 | Decrease |
| Vermis VI-VII | 47.0–49.4; 52.6–55.1; 58.3–90.0 | -2.61; -2.47; -7.27 | Decrease |
| Vermis VIII | 70.7–87.4 | -2.35 | Decrease |
| Vermis IX | 57.5–90.0 | -2.32 | Decrease |
| Vermis VI-IX | 46.1–90.0 | -12.11 | Decrease |
| Vermis X | 70.5–85.9 | -0.91 | Decrease |
| Bilateral Anterior Lobe | 62.9–90.0 | -50.48 | Decrease |
| Left Anterior Lobe | 62.6–90.0 | -13.44 | Decrease |
| Right Anterior Lobe | 63.0–90.0 | -35.88 | Decrease |
| Bilateral Posterior Lobe | 36.0–90.0 | -531.20 | Decrease |
| Left Posterior Lobe | 36.0–90.0 | -300.25 | Decrease |
| Right Posterior Lobe | 36.0–90.0 | -247.13 | Decrease |
| Bilateral Flocculonodular Lobe | 69.7–86.4 | -1.57 | Decrease |
| Whole Vermis | 42.0–90.0 | -12.92 | Decrease |
| Left Cerebellar Hemisphere | 36.0–90.0 | -299.46 | Decrease |
| Right Cerebellar Hemisphere | 36.0–90.0 | -261.84 | Decrease |
| Corpus Medullare | 52.1–90.0 | -79.46 | Decrease |
| Total Cerebellum | 36.0–90.0 | -731.20 | Decrease |
| *Note.* Significant age ranges were identified by evaluating the first derivative of the fitted smooth age effect at 1,000 evenly spaced points across the observed range. Ages where the simultaneous 95% confidence interval excluded zero indicate a significant instantaneous rate of volume change. The peak rate reflects the maximum instantaneous rate of volume change for each significant age window. Regions that have multiple significant ranges are separated by semicolons. n.s. = no significant change. | | | |

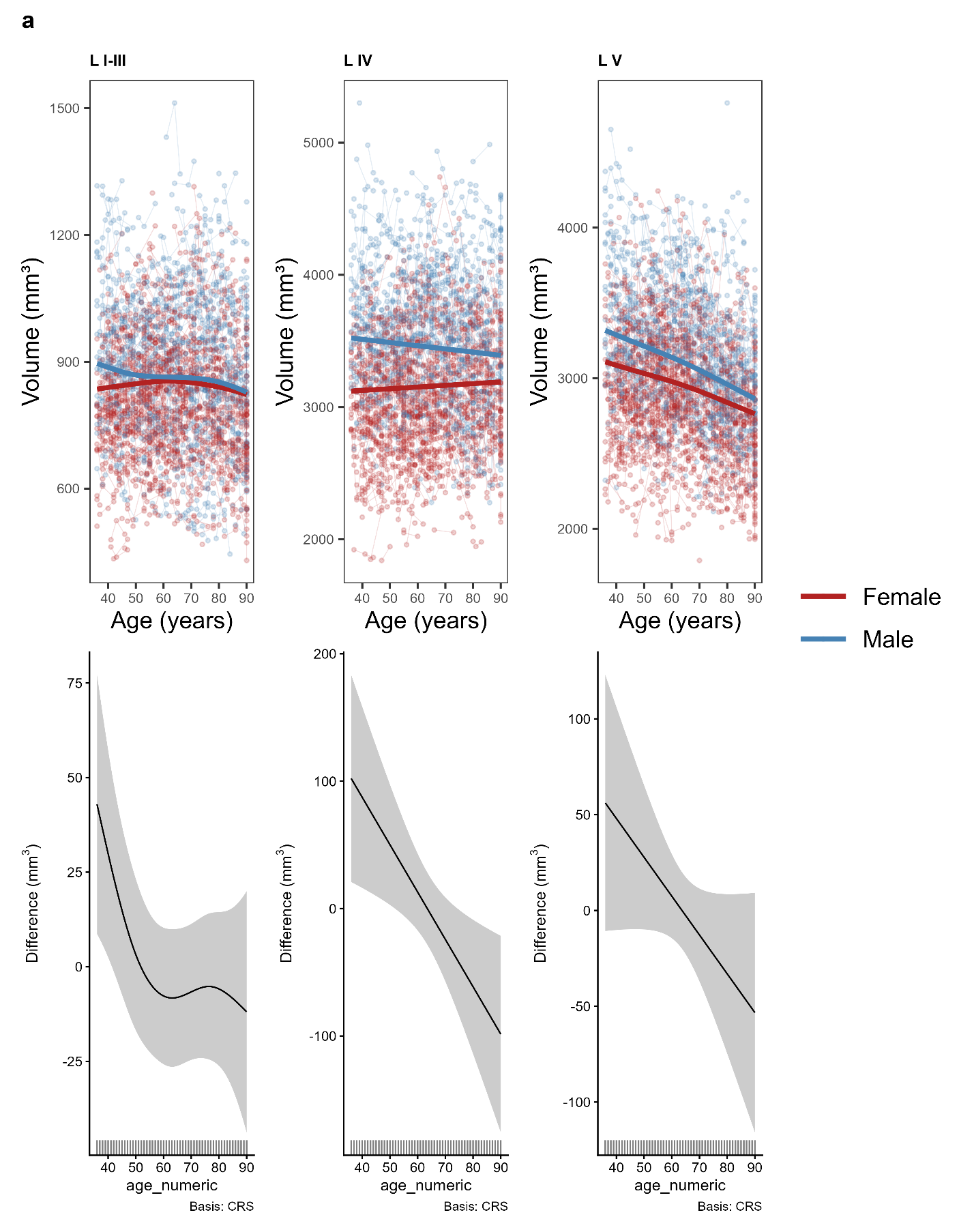

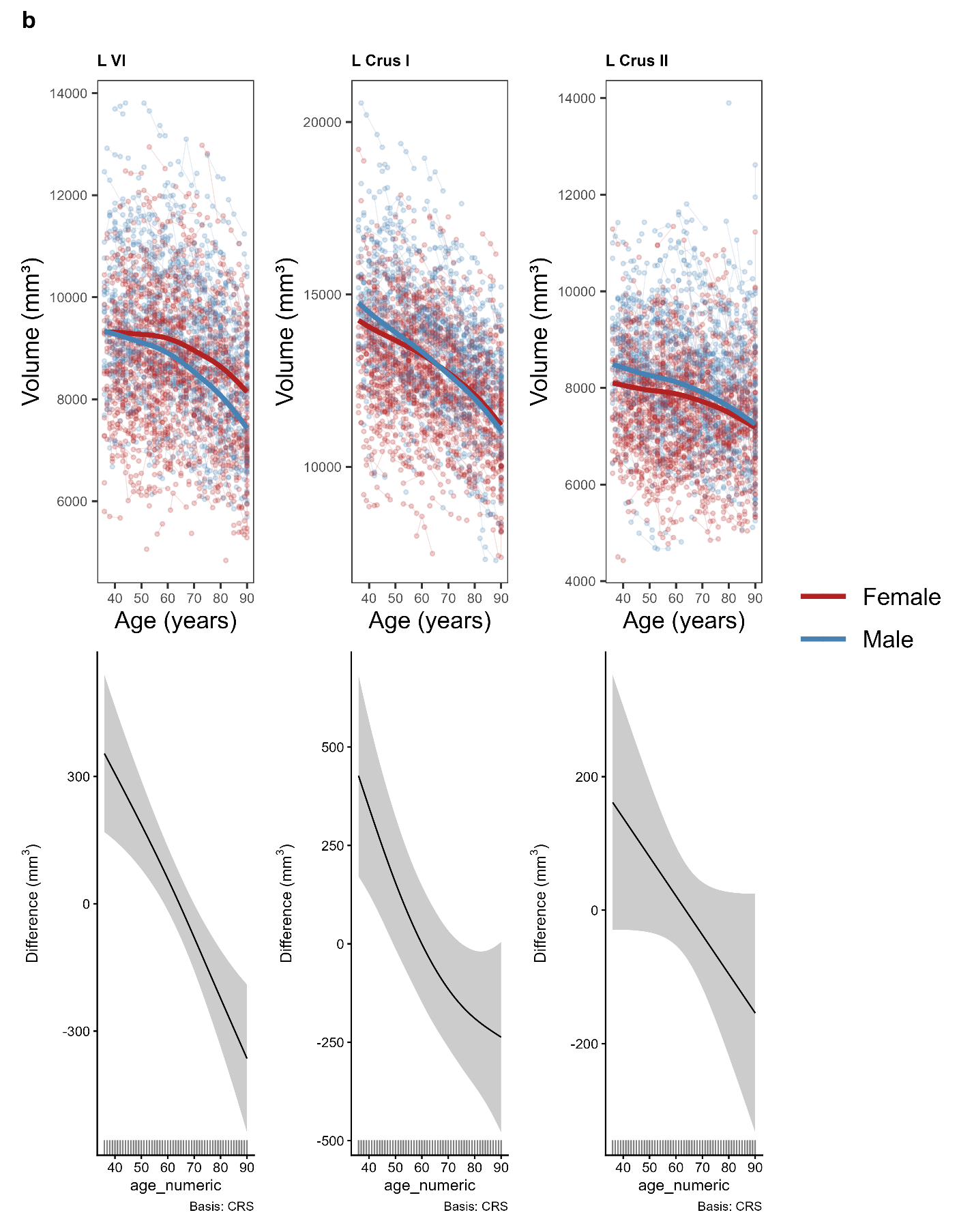

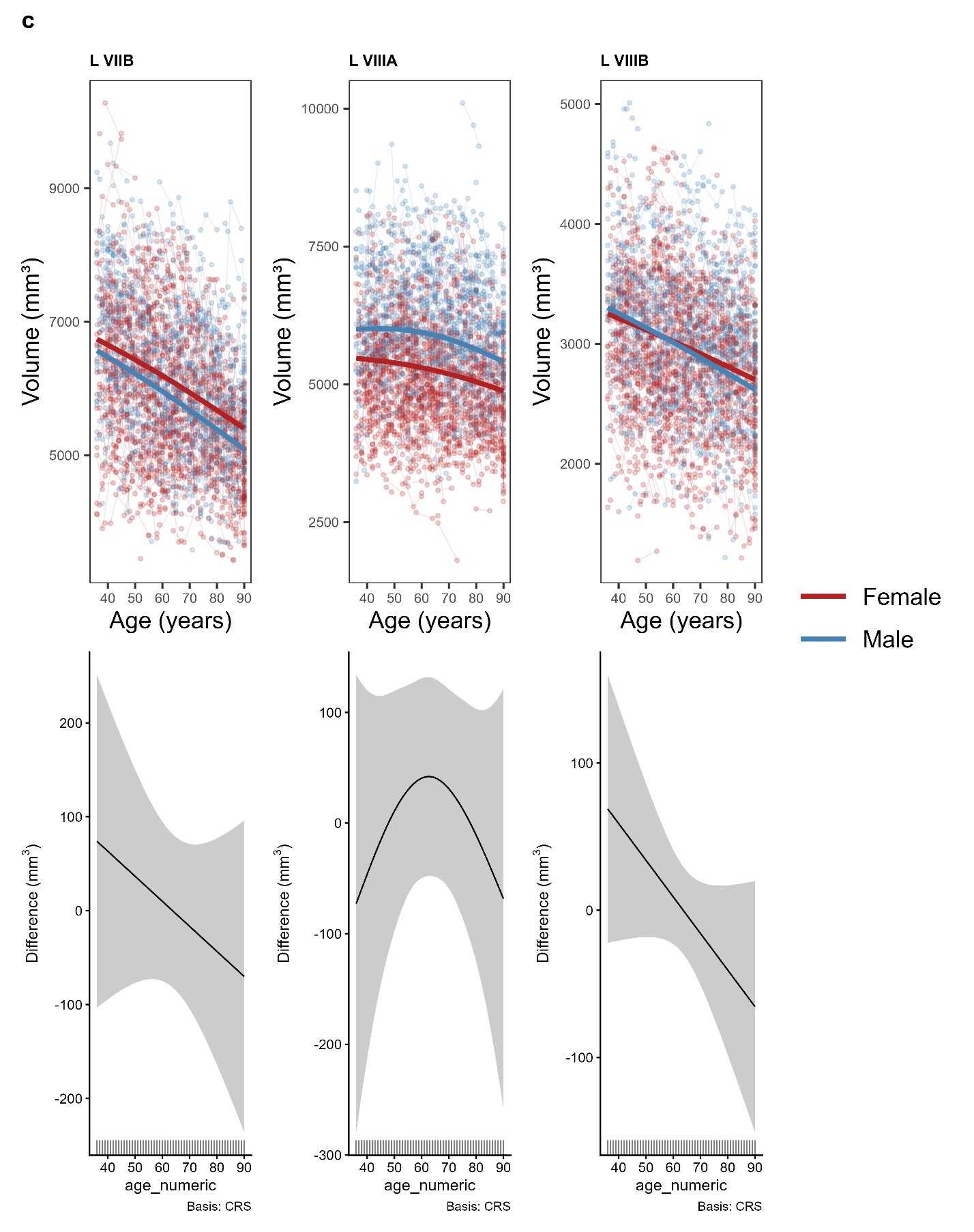

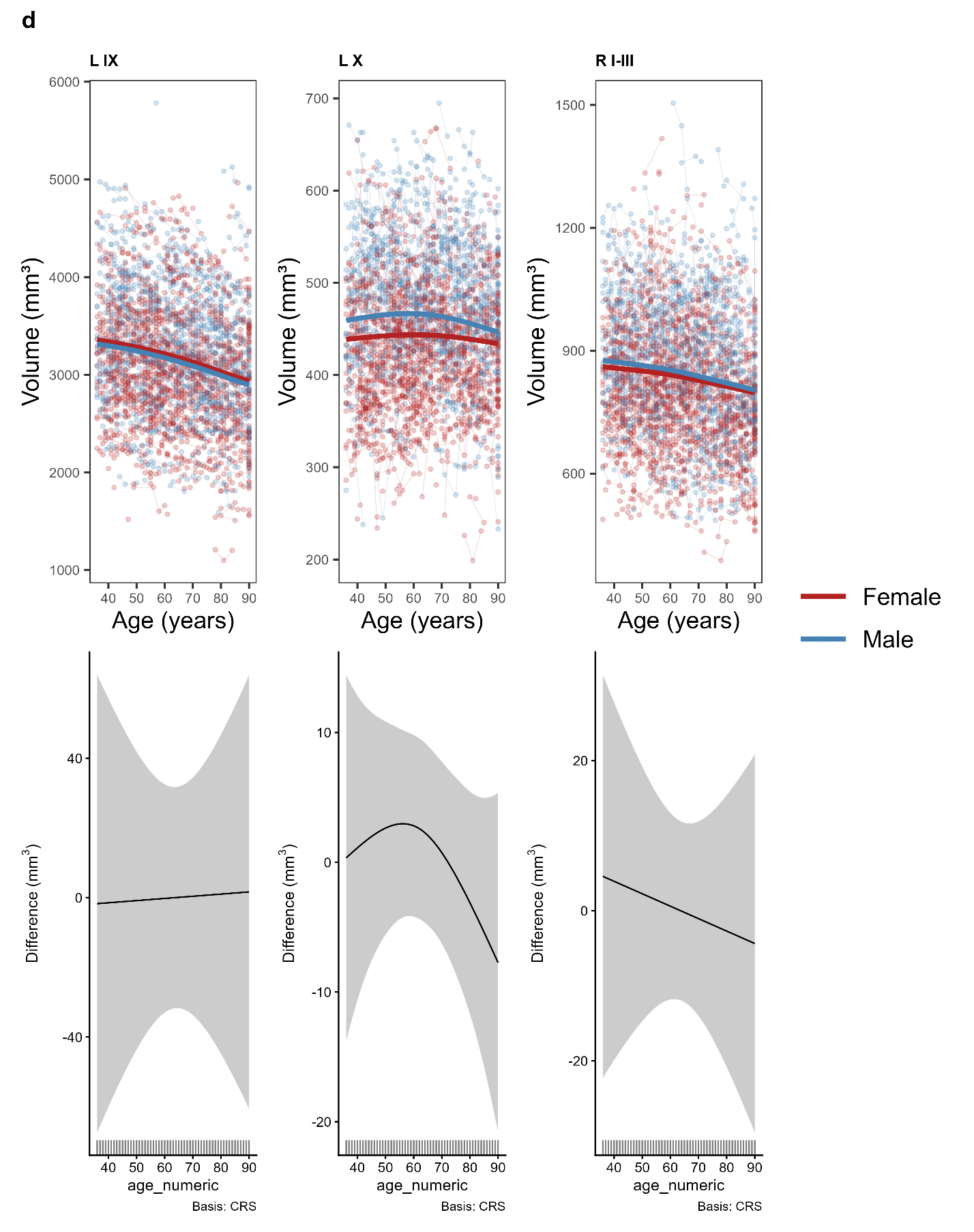

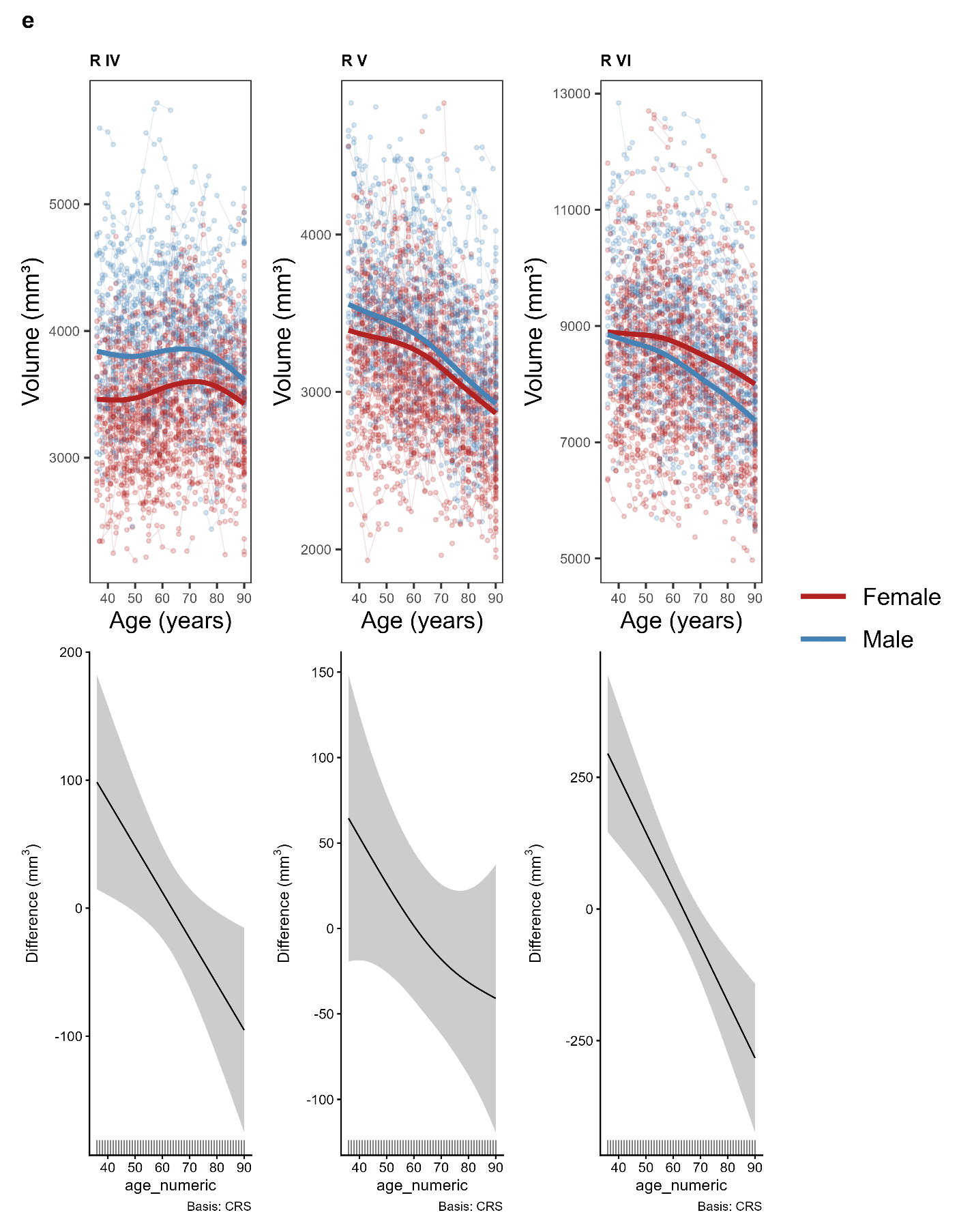

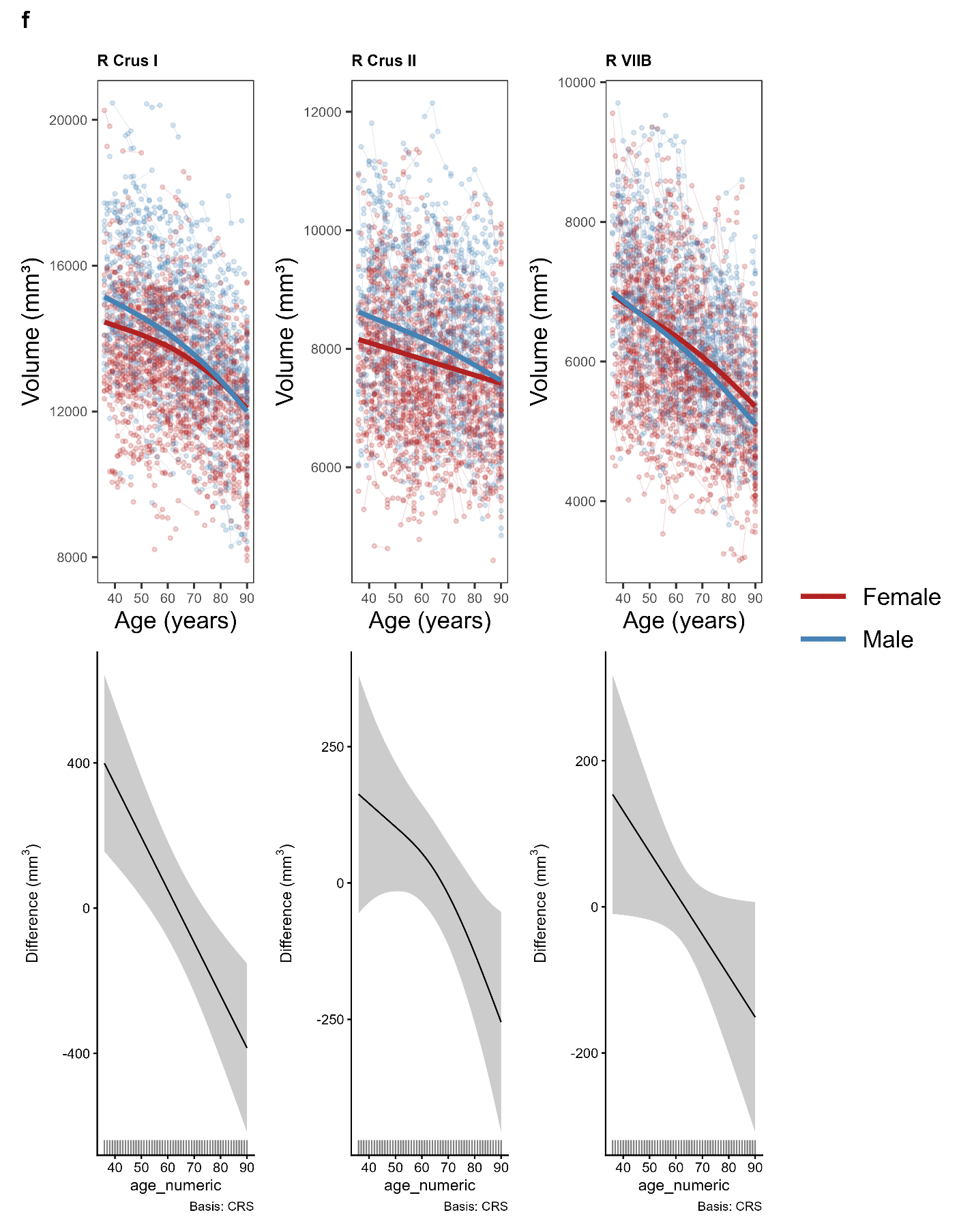

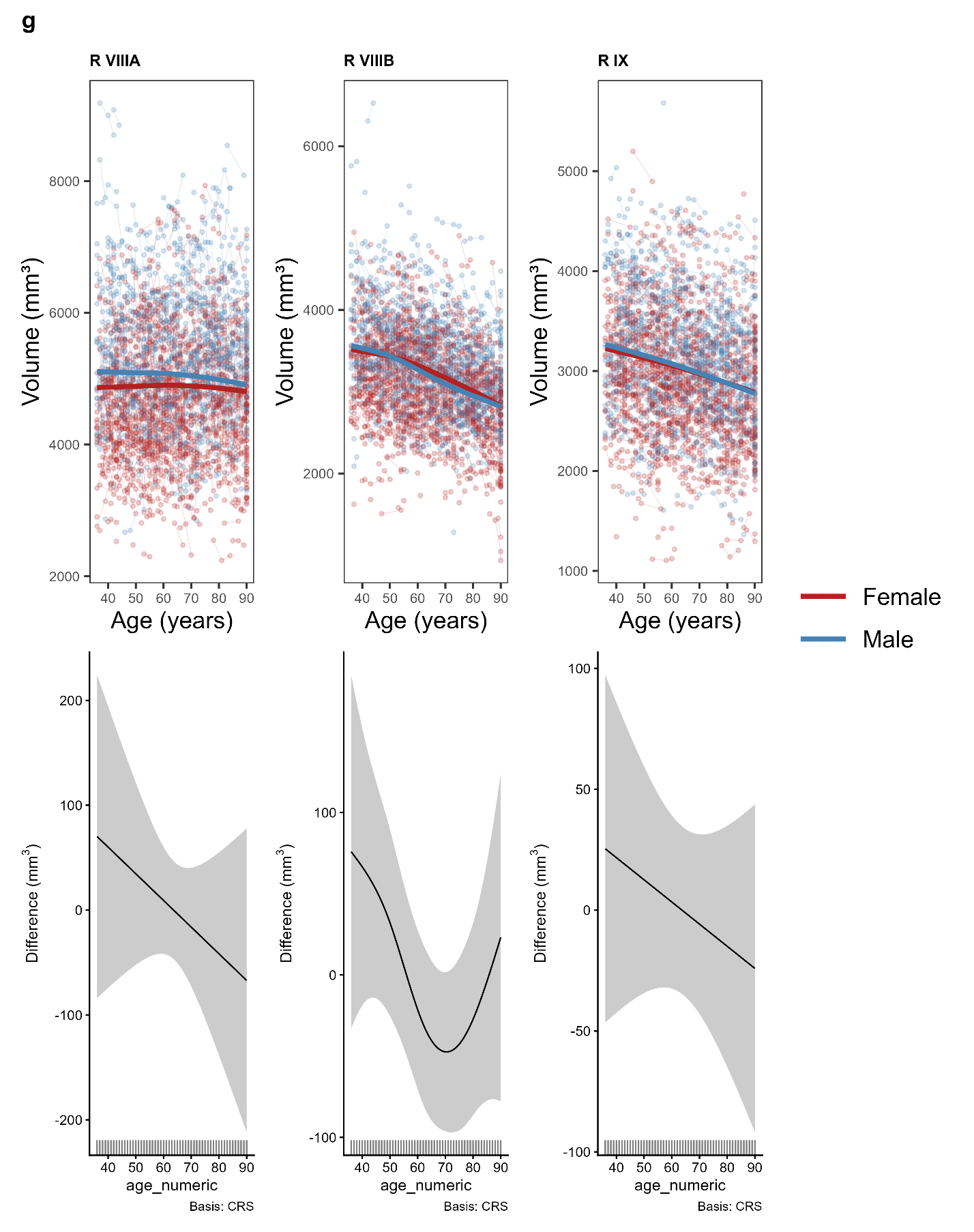

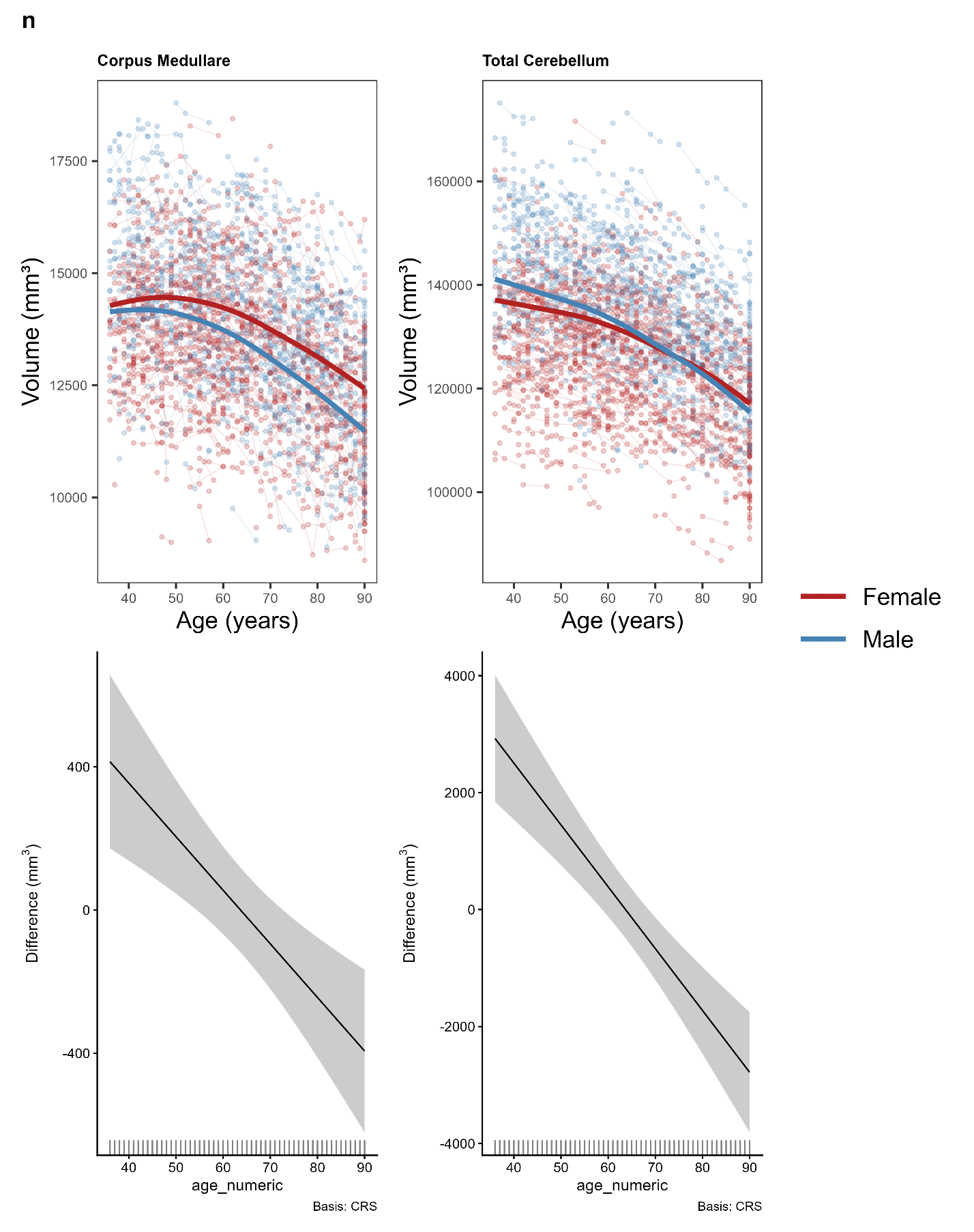

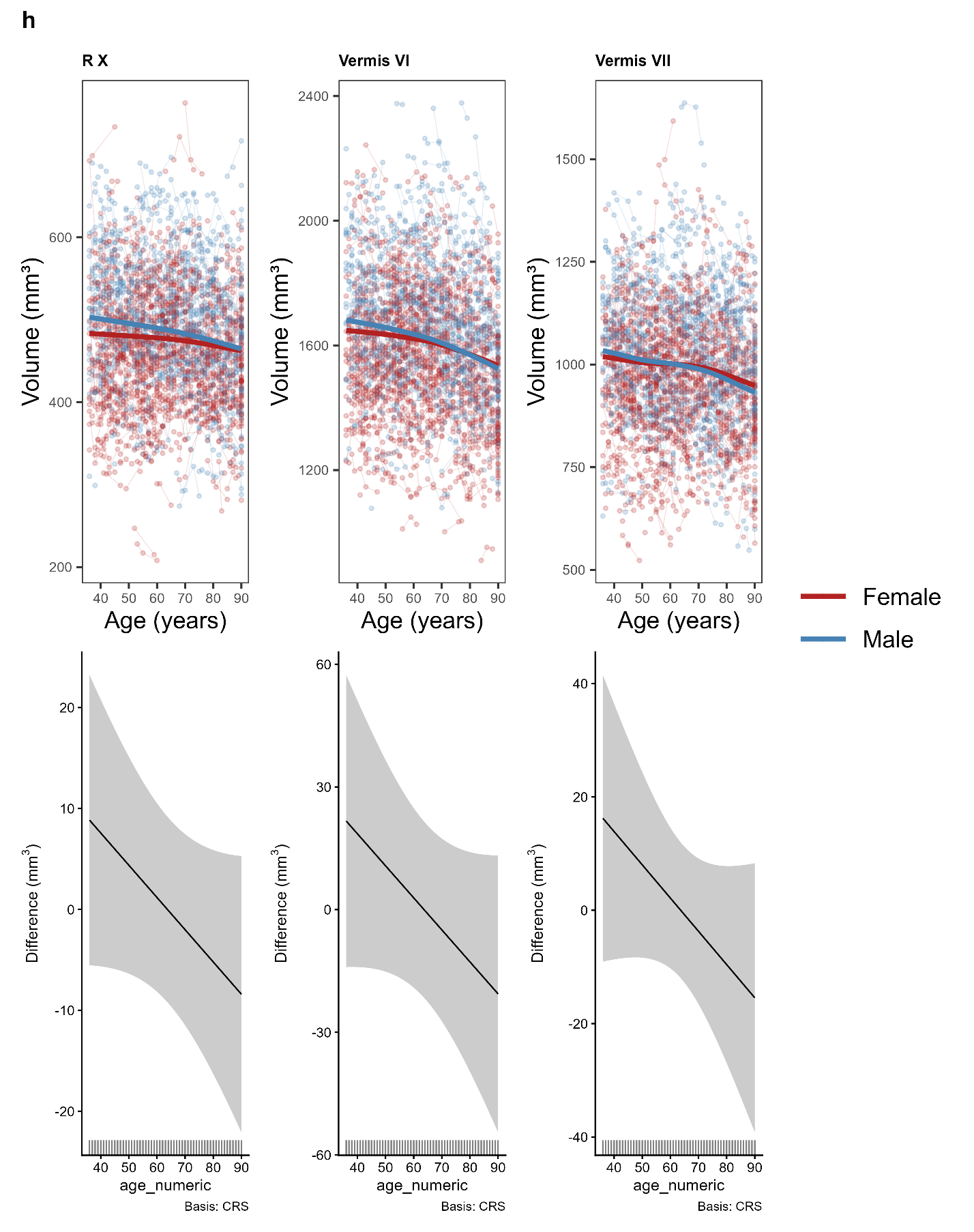

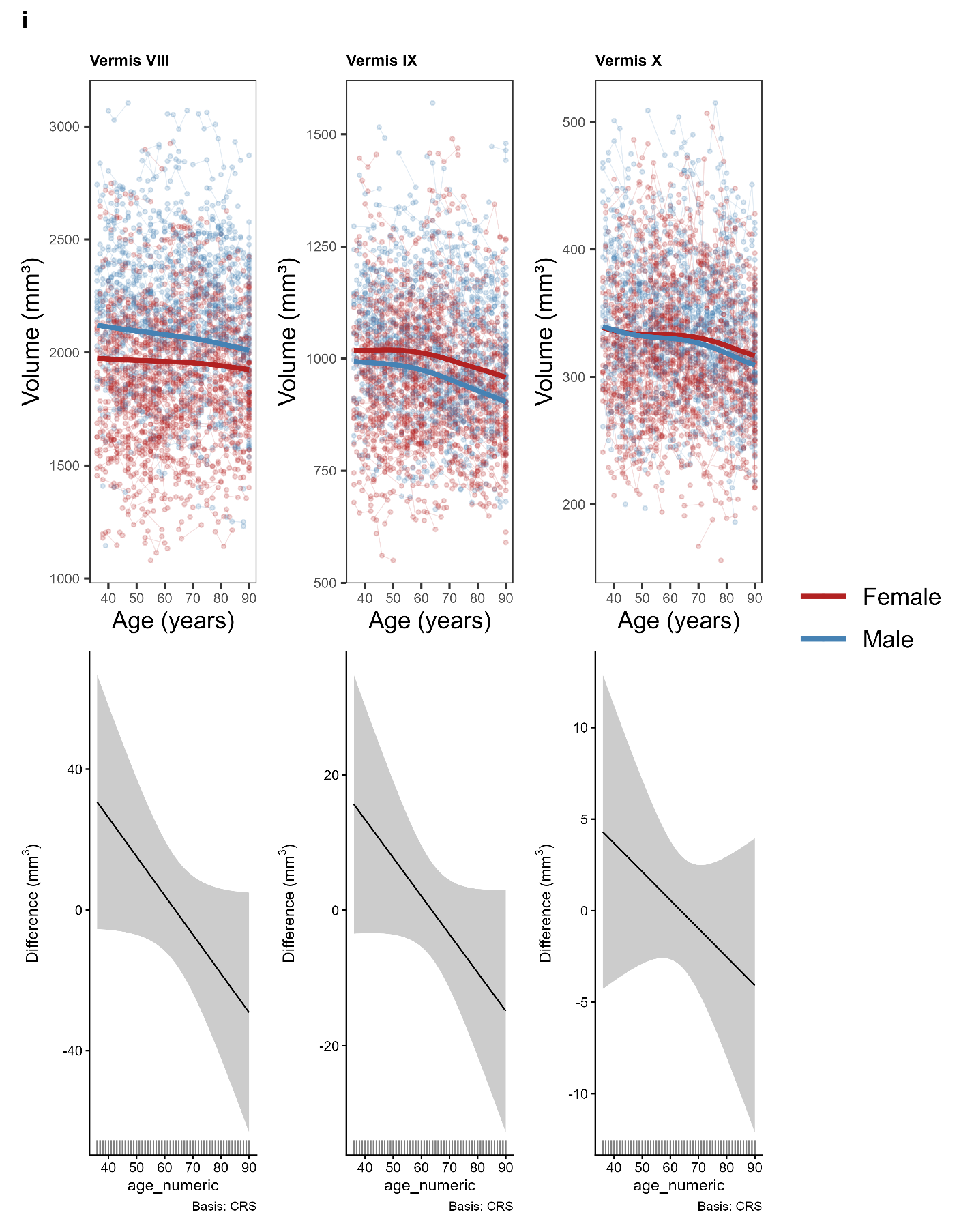

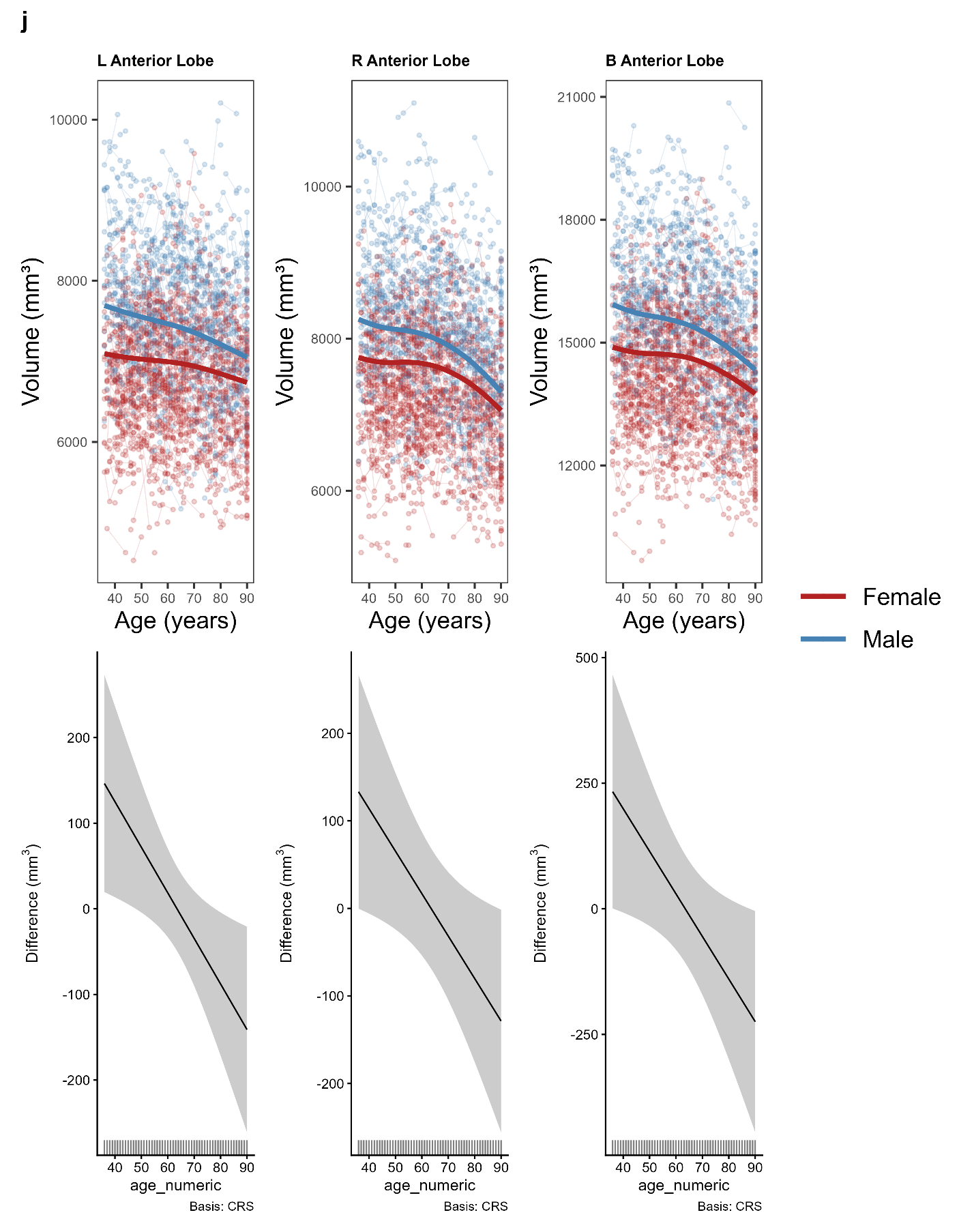

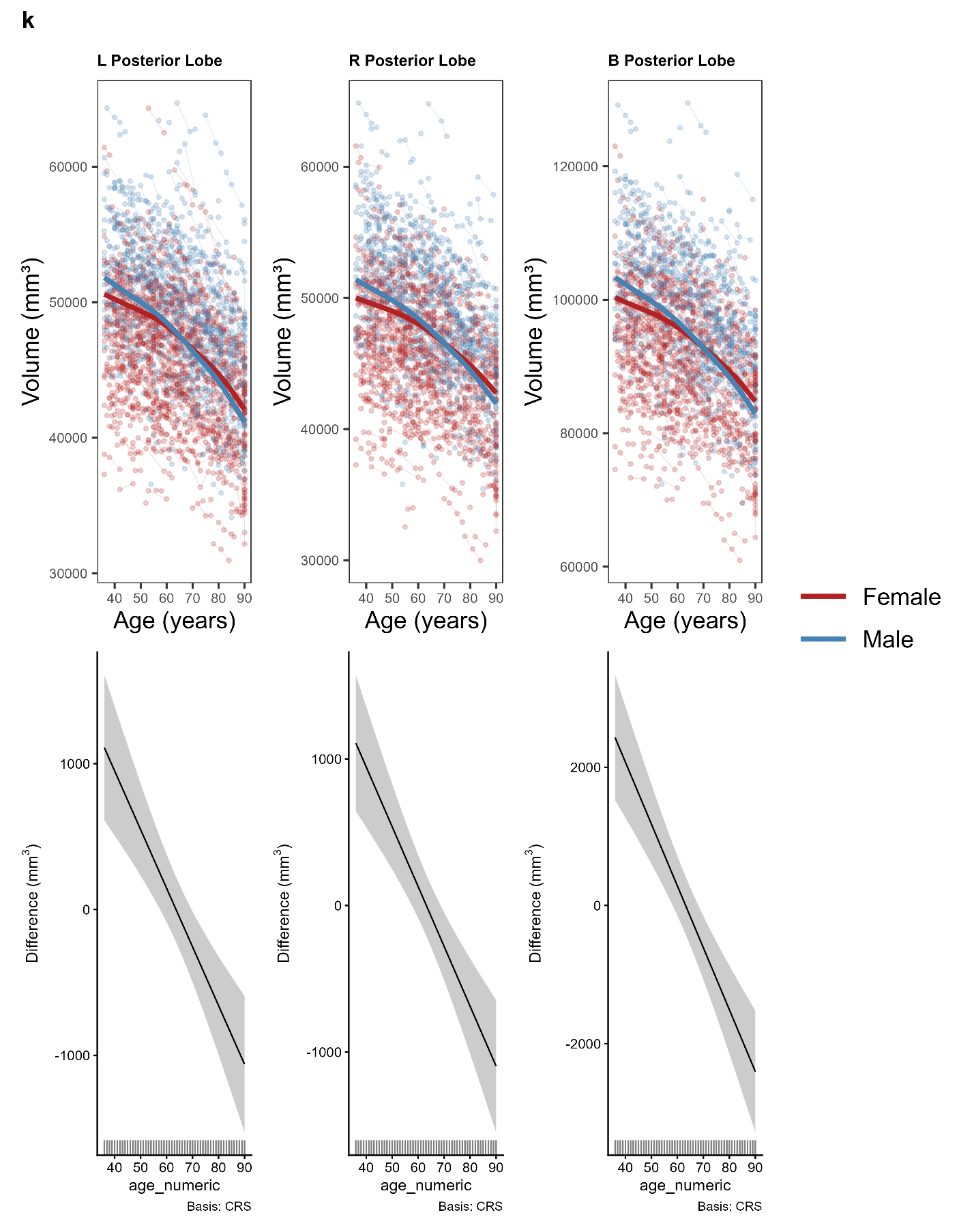

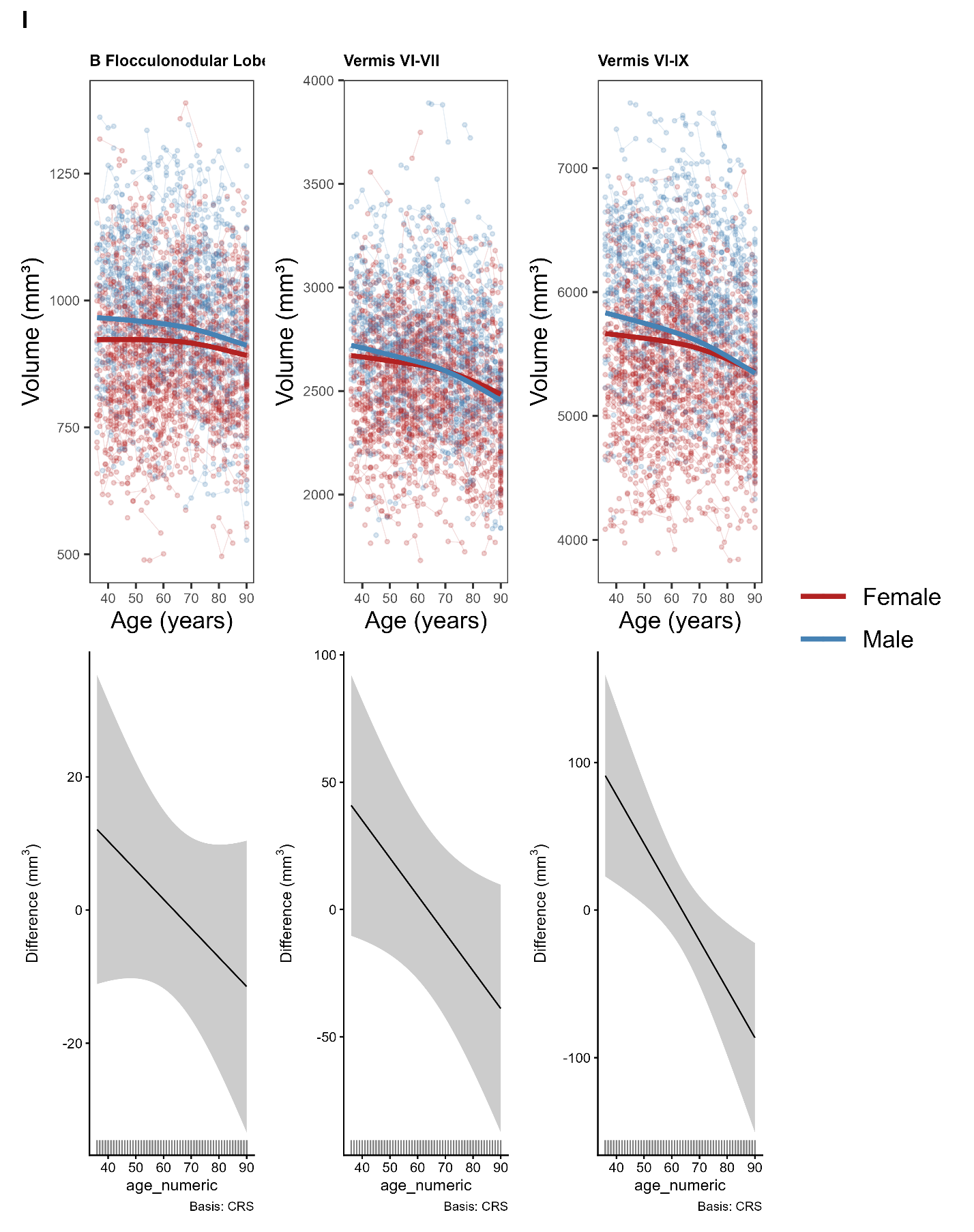

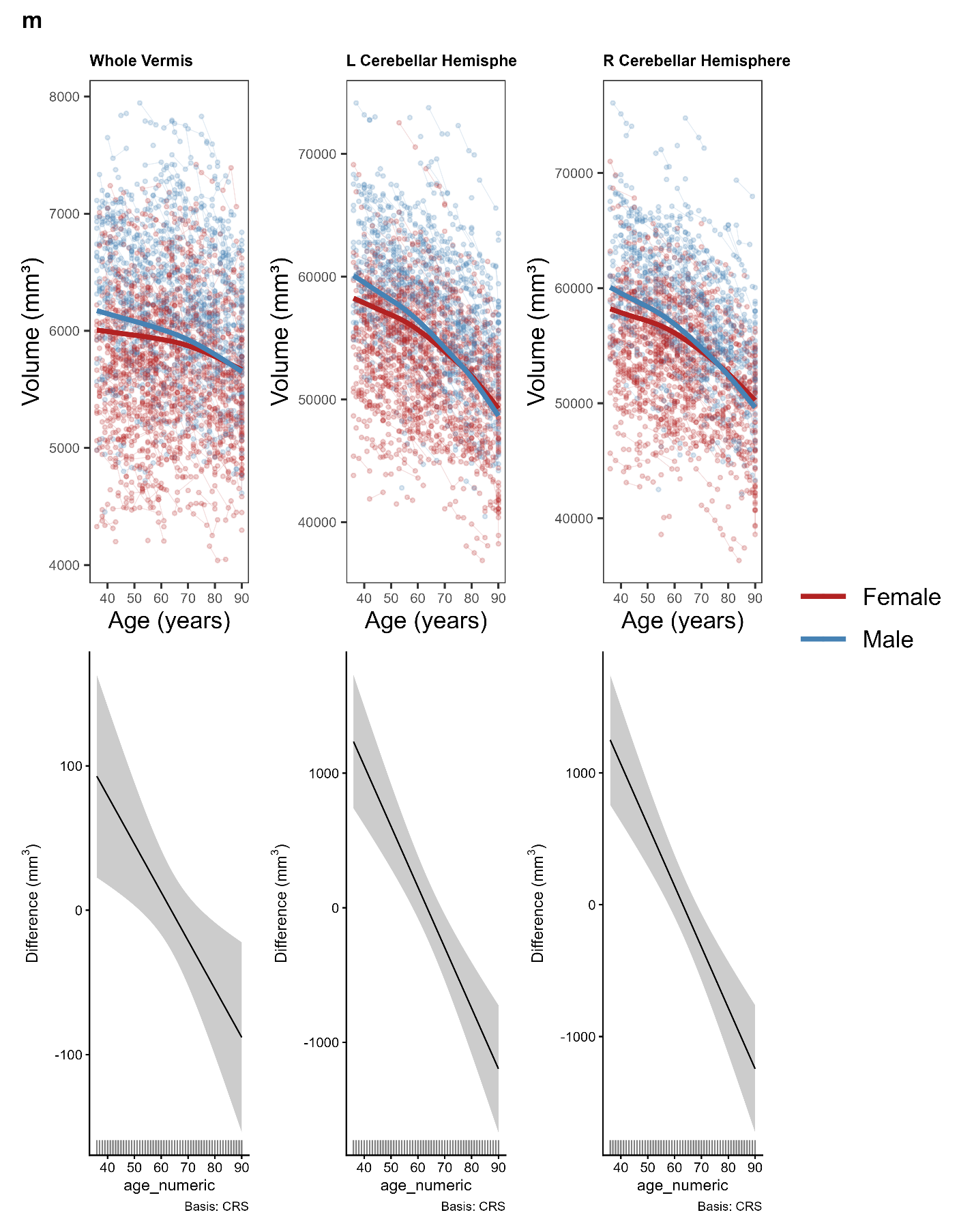

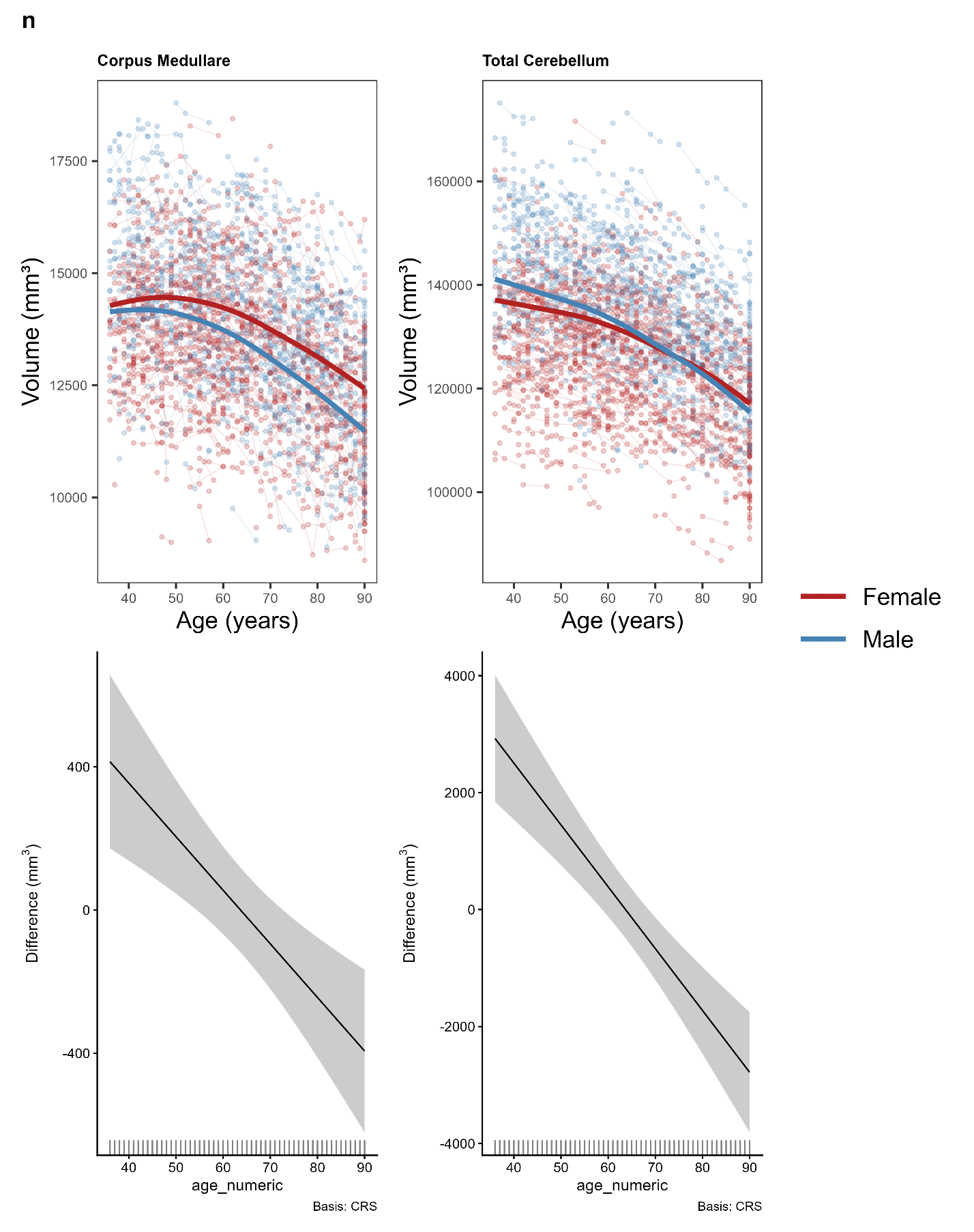

**Supplemental Fig. 4.** Top rows (**Supplemental** **Fig. 4a** – **Supplemental** **Fig. 4n**) shows predicted model trajectories for males (blue) and females (red) from the sex*age interaction model overlaid on uncorrected volumetric data across visits for each ROI. Bottom rows (**Supplemental** **Fig. 4a** – **Supplemental** **Fig. 4n**) show partial effect plots illustrating how volumetric trends (mm³) may deviate for mean from females (reference) across age. Values at zero indicate no difference; positive values are indicative of greater male volume at the respective age, and vice versa for negative values. Each partial effect plot was derived from the sex*age interaction model, paired with 95% confidence interval grey bands. Smooth interaction terms used cubic regression splines as the basis function (Basis: CRS). Linear deviations illustrate a linear deviation from reference (i.e., edf = 1.00) while nonlinear effects indicate nonlinear deviations (i.e., edf = 2.00). Models controlled for eTIV, and random subject and scanner effects. Abbreviations in the plot titles are as follows: R = Right, L = Left, B = Bilateral.

**Supplemental Table 2.** Comparison of model statistics across cerebellar ROIs between primary and sensitivity analyses.

|  |  |  | Primary Model | | | Sensitivity Model | | |  |
| --- | --- | --- | --- | --- | --- | --- | --- | --- | --- |
| **Model** | **Sensitivity Analysis** | **ROI** | **edf** | ***F*** | ***p*** | **edf** | ***F*** | ***p*** | **Change (+/-)** |
| Cross-sectional Model | Blood Pressure | Left I-III | 2.28 | 2.56 | 0.045 | 2.56 | 2.07 | 0.054 | - |
|  | Race | Left I-III | 2.28 | 2.56 | 0.045 | 2.24 | 1.91 | 0.10 | - |
|  | Race | Vermis VII | 1.64 | 3.05 | 0.045 | 1.22 | 2.61 | 0.07 | - |
|  | Race | Vermis VIII | 1.01 | 1.92 | 0.17 | 1.21 | 12.88 | <0.01 | + |
|  | Ethnicity | Left X | 2.37 | 2.44 | 0.07 | 2.38 | 2.78 | 0.04 | + |
|  | Education | Left I-III | 2.28 | 2.56 | 0.045 | 2.27 | 2.47 | 0.051 | - |
|  | MoCA | Bilateral Flocculonodular Lobe | 2.33 | 4.51 | <0.01 | 2.03 | 2.15 | 0.10 | - |
| Longitudinal Age Model | Race | Right VIIIA | 2.17 | 1.99 | 0.11 | 2.08 | 4.21 | 0.01 | + |
|  | Education | Vermis VIII | 2.69 | 7.56 | <0.01 | 2.27 | 1.55 | 0.16 | - |
|  | MoCA | Left X | 2.87 | 3.84 | <0.01 | 2.52 | 2.27 | 0.09 | - |
| Sex*Age Interaction Model | Race | Left I-III | 2.67 | 3.46 | 0.06 | 2.69 | 3.85 | 0.04 | + |
|  | Race | Left Crus II | 1.00 | 3.27 | 0.07 | 1.00 | 4.10 | 0.045 | + |
|  | Race | Vermis VI-VII | 1.00 | 4.22 | 0.04 | 1.00 | 2.92 | 0.09 | - |
|  | Race | Right VIIB | 1.00 | 3.89 | 0.049 | 1.00 | 2.93 | 0.09 | - |
|  | Ethnicity | Vermis VI-VII | 1.00 | 4.22 | 0.04 | 1.00 | 3.80 | 0.052 | - |
|  | Ethnicity | Right VIIB | 1.00 | 3.89 | 0.049 | 1.00 | 3.72 | 0.054 | - |
|  | Education | Right Crus II | 1.77 | 4.53 | 0.04 | 1.54 | 4.46 | 0.053 | - |
|  | Education | Right VIIB | 1.00 | 3.89 | 0.049 | 1.00 | 2.95 | 0.09 | - |
|  | MoCA | Right Crus II | 1.77 | 4.53 | 0.04 | 1.65 | 4.48 | 0.050 | - |
|  | MoCA | Right VIIB | 1.00 | 3.89 | 0.049 | 1.00 | 3.79 | 0.052 | - |
|  | MoCA | Vermis VIII | 1.00 | 3.42 | 0.06 | 1.00 | 3.94 | 0.047 | + |

*Note.* The change column reflects whether the sensitivity analysis saw a gain (+) or loss (-) of significance (*p* < 0.05) for the smooth age effect of the cross-sectional and longitudinal models, and sex*age interaction for the interaction model. If an ROI or analysis was not listed for a given model, that signifies no change in main effects occurred. edf = Effective Degrees of Freedom.
